# Pandemic preparedness through genomic surveillance: Overview of mutations in SARS-CoV-2 over the course of COVID-19 outbreak

**DOI:** 10.1101/2023.08.12.553079

**Authors:** Fares Z. Najar, Chelsea L. Murphy, Evan Linde, Veniamin A. Borin, Huan Wang, Shozeb Haider, Pratul K. Agarwal

**Affiliations:** High-Performance Computing Center, Oklahoma State University, Stillwater, Oklahoma; Department of Physiological Sciences, Oklahoma State University, Stillwater, Oklahoma; University College London School of Pharmacy, Pharmaceutical and Biological Chemistry, London, United Kingdom; University College London Centre for Advanced Research Computing, London, United Kingdom

## Abstract

Genomic surveillance is a vital strategy for preparedness against the spread of infectious diseases and to aid in development of new treatments. In an unprecedented effort, millions of samples from COVID-19 patients have been sequenced worldwide for SARS-CoV-2. Using more than 8 million sequences that are currently available in GenBank’s SARS-CoV-2 database, we report a comprehensive overview of mutations in all 26 proteins and open reading frames (ORFs) from the virus. The results indicate that the spike protein, NSP6, nucleocapsid protein, envelope protein and ORF7b have shown the highest mutational propensities so far (in that order). In particular, the spike protein has shown rapid acceleration in mutations in the post-vaccination period. Monitoring the rate of non-synonymous mutations (K_a_) provides a fairly reliable signal for genomic surveillance, successfully predicting surges in 2022. Further, the external proteins (spike, membrane, envelope, and nucleocapsid proteins) show a significant number of mutations compared to the NSPs. Interestingly, these four proteins showed significant changes in K_a_ typically 2 to 4 weeks before the increase in number of human infections (“surges”). Therefore, our analysis provides real time surveillance of mutations of SARS-CoV-2, accessible through the project website http://pandemics.okstate.edu/covid19/. Based on ongoing mutation trends of the virus, predictions of what proteins are likely to mutate next are also made possible by our approach. The proposed framework is general and is thus applicable to other pathogens. The approach is fully automated and provides the needed genomic surveillance to address a fast-moving pandemic such as COVID-19.

## INTRODUCTION

COVID-19, caused by the SARS-CoV-2 virus, ground the entire world to a halt in a matter of a few weeks. It aptly revealed that an airborne virus with the ability to quickly mutate, combined with high levels of transmission and fatality, can rapidly cause a world-wide pandemic. The response from the medical and research community was swift, leading to the development of mRNA-based vaccines and several antiviral drugs. Unfortunately, even after three years of this worldwide pandemic, the situation remains alarming. New variants have emerged and continue to emerge, causing new waves of infections and fatality. Our ability to respond to the future pandemics, will depend on our ability to stay ahead of the pathogens. Therefore, there is wide interest in approaches for genomic surveillance as a means for pandemic preparedness and response. To this end, several million SARS-CoV-2 sequences have already been obtained in a hitherto unprecedented global effort. These sequences are being deposited by various health labs in public repositories such as the NCBI’s GenBank and GISAID.

SARS-CoV-2 was first sequenced in 2020 from a sample in Wuhan, China. This sequence, widely known as the “Wuhan” sequence,^1^ showed that SARS-CoV-2 is a positive sense RNA virus that encodes 26 proteins (Figure 1). The expressed proteins include: four structural proteins known as the envelope, the membrane protein, nucleocapsid protein, and the spike protein;^2^ 16 non-structural proteins (NSP 1–16); and six accessory proteins as open reading frames (ORFs), 3a, 6, 7a, 7b, 8, and 10.^3^ The two large overlapping open read frames, ORF1a and ORF1b, are transcribed and later split into different NSPs. The overlapping occurs at a frameshift to allow the continuous transcription of NSP12.^1^

**Figure 1:**
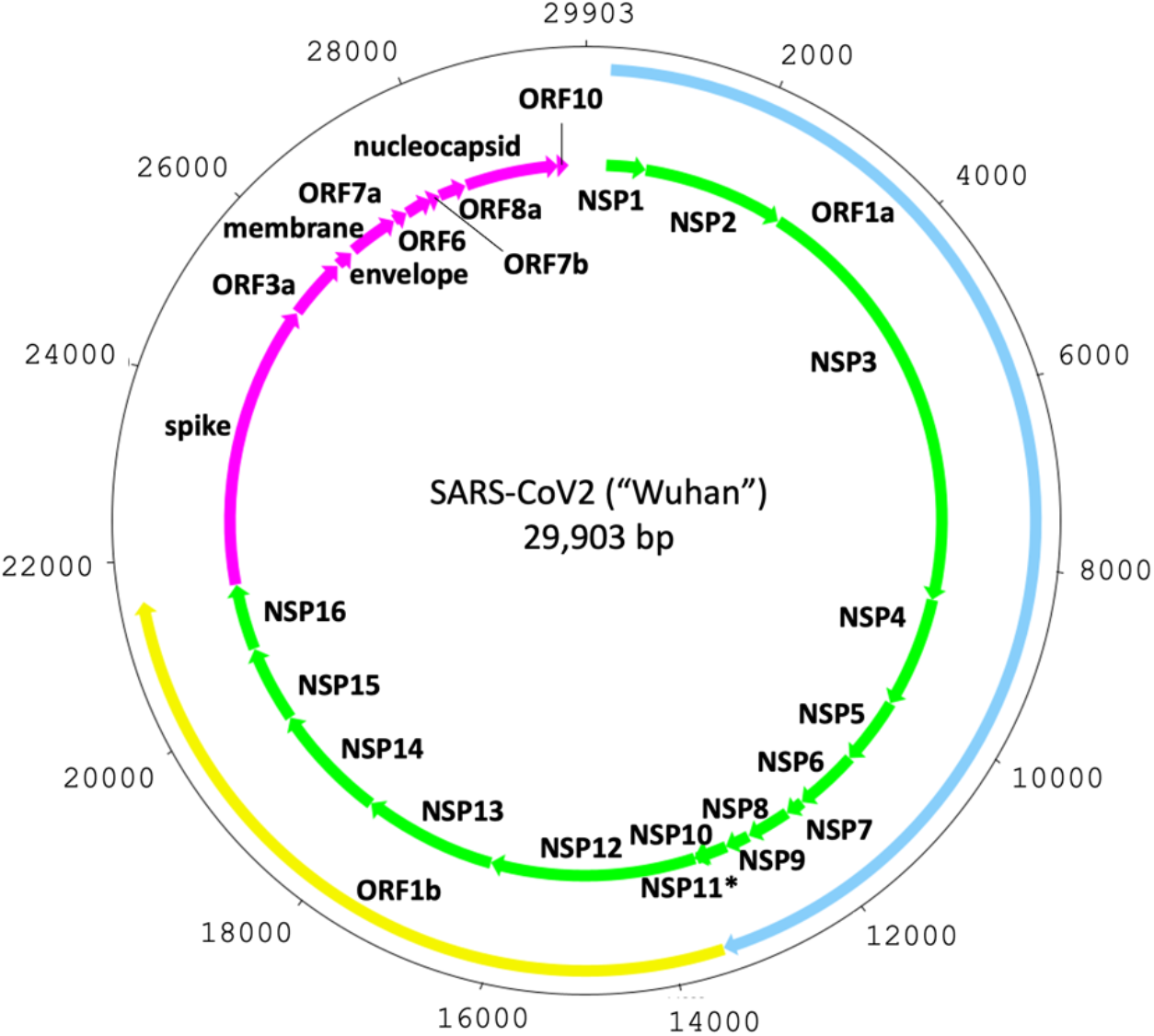
A schematic overview of the SARS-CoV-2 genome. Information based on the “Wuhan” reference sequence. The non-structural genes are shown in green while the structural genes are marked in magenta. The ORF1a and ORF1b boundaries are shown in light blue and yellow respectively. The position of NSP11 (marked by *), which is too small to be visible at this schematic’s scale, is not shown.

For the genomic surveillance to be useful for pandemic (such as COVID-19) preparedness, several aspects are required. First, the SARS-CoV-2 proteins that are rapidly changing need to be identified in real-time (or near real-time). Ideally, this information also needs to be available for different geographic locations separately, as the outbreaks show different regional trends. Information on how different variants are mutating is also important, as at any given time there can be (and have been) different variants of the virus in the host population. Collectively, this set of information should enable prediction of what mutated versions of the protein will be presented in the next variants of concern (VOC). However, for this information to be useful beyond the research community, such as aiding the medical community in making practical but critical decisions, the genomic surveillance should be able to clearly predict future outbreaks or increase in number of cases of infections (“surges”). Furthermore, in order to make an impact on mitigation strategies, it is vital that all of this analysis needs to be performed in real-time.

From a molecular perspective, a detailed analysis of mutations in the virus proteins are needed for understanding of the new outbreaks. Number and types of mutations in individual proteins associated with various variants could shed light on severity of the ongoing or future outbreaks. This information could also help in designing new vaccines and drugs, and combat antiviral resistance which could arise from new mutations. Furthermore, the effectiveness of the antibody-based viral detection methods is also prone to large mutational changes in the different proteins. (Note that some of this information has been generated in-house by vaccine and drug developing groups; however, access to such information remains lacking for the broader scientific community at large.) A detailed understanding of mutational propensities of various proteins would be the key to pre-emptively designing antibodies that will be able to detect mutated proteins, as well as vaccines that are effective against new VOCs. Similarly, relating mutations, molecular structure, and functional mechanism could also provide new opportunities for effective antiviral drugs.

Recently, we described our real-time genomic surveillance of SARS-CoV-2 based on mutation analysis of viral proteins as a methodology for *a priori* determination of surge in number of infection cases. The results are available at http://pandemics.okstate.edu/covid19/, and are updated daily as new virus sequences become available. At present, over 8 million SARS-CoV-2 genome sequences collected all over the world are available from GenBank (https://www.ncbi.nlm.nih.gov/sars-cov-2/) and were used for analysis of 26 SARS-CoV-2 genes. As a result of this effort we were able to predict the upcoming Omicron BA.5 related surge in end of June 2022, which was confirmed in the upcoming weeks of July 2022. We also issued a warning in September 2022, which was confirmed by infections increase in Europe and several individual European countries. Furthermore, a new watch was issued in the first week of January 2023, which corresponded to the current late January 2023 surge.

Here, we describe the detailed mutations overview of all the 26 proteins and ORFs of SARS-CoV-2 over the course of the COVID-19 outbreak. Our analysis is based on the unprecedented SARS-CoV-2 sequencing campaign undertaken by the medical community and public health labs worldwide. Furthermore, the information available for the number of daily infection cases from public databases has allowed us to make real-time prediction about impending surges in infections. Our approach is model-free, which allows for easy interpretation by the research community as well as the medical community and public health agencies for preparedness purposes. Based on the results, we have also developed an approach to predicting which heavily mutated proteins will likely be present in the upcoming variants. The presented approach and framework is general and can be applied to other future infectious diseases.

## METHODS

The raw SARS-CoV-2 genomic sequences data and the number of COVID-19 infection cases are continually obtained from the sources described below. The genomic sequences are first carefully filtered for quality control, and sequences passing quality control are used for identification of mutations and calculations of non-synonymous and synonymous mutation rates and their ratio for each of the listed 26 proteins separately.

### Data and data sources

Data for the number of reported COVID-19 cases is accessed from Our World In Data project (https://ourworldindata.org/coronavirus-source-data).^4^

### Genomic sequence data

An in-house pipeline of scripts (using Linux commands) was designed around the eUtils tools^5^ from NCBI in order to download and process the SARS-CoV-2 records from NCBI’s GenBank (https://www.ncbi.nlm.nih.gov/genbank/). Briefly, we used *esearch* and *efetch* commands to obtain these GenBank records. The search string “SARS-CoV-2”, refined to “SARS-CoV-2 [ORGN]”, was used to download the identified records in the GenBank text format. After workflow optimization post May 2022, the search process used NCBI’s newer *datasets* and *dataformat* command-line tools to identify sequences of interest, while continuing to use the *efetch* tool to download records in the GenBank text format. Collectively, a total of 8,026,343 records were searched, and a total of 3,596,381 sequences matching the search criterion were downloaded and used as of July 20^th^, 2023.

### Quality control

Incomplete and ambiguous SARS-CoV-2 genomic sequences and records containing incomplete collection dates were filtered out using the designed pipeline. For the records passing the quality control steps, the nucleotide sequence for each gene was extracted. A non-redundant set of the extracted nucleotide sequences was derived and translated to the cognate amino acid sequences. In the final phase of the pipeline, the accession numbers for each viral isolate, the nucleotide sequences, the associated protein sequences, the collection dates, and the country of collection were stored in an internal SQLite relational database, where they were indexed with unique identifiers to allow for the retrieval and analysis of any part of the parsed data.

### Frequency of data updates

As of April 2022, the described sources are being monitored daily for updates. New data is continually downloaded and used for analysis through automated pipelines.

### Amino acid alignments and mutational propensity

For each protein, we used Clustal-Omega^6^ to align the amino acids from each variant. The total number amino acid changes (substitutions and deletions) were calculated and summed up for each gene. Those values were divided by the total amino acid length for each gene to obtain the mutational rank, where higher mutational ranks correspond to a higher propensity for mutations for that gene.

### Alignments and non-synonymous (K_a_), synonymous (K_s_) and K_a_/K_s_ ratio calculations

The translated protein and nucleotide sequences were aligned using *clustal-omega*^6^ and *Pal2Nal*^7^ programs to align the codons with their associated amino acids. The resulting alignments were then processed through the program *kaks_calculator*^8^ to calculate non-synonymous (K_a_) and synonymous (K_s_) values and their ratio K_a_/K_s_ values which were used to assess the mutational adaptation for each protein. The parameters required for the *kaks_calculator* program were based on the maximum-likelihood method derived from the work of Goldman and Yang.^9^ The first reported SARS-CoV-2 genomic sequence (the “Wuhan” sequence) was used as a reference for all the K_a_, K_s_ and K_a_/K_s_ calculations. While we explored the possibility of using other sequence(s) as references (e.g., the previous day or the previous month), the increasing number of variations available each day makes it prohibitive to select a representative consensus sequence on an ongoing basis. Additionally, we found that using the Wuhan sequence as a reference provided the most intuitive and interpretable results. We observed the mean K_a_, K_s_, and K_a_/K_s_ over the entire documented timeline of COVID-19 cases to discover if there is a correlation between these parameters’ measurements and the infection surges, and to help reveal the genotypic and virulence natures of the variants of interest preceding the surges.

The targeting of both the non-synonymous and synonymous mutation values was selected due to its ability to observe changes at both the protein and the nucleotide levels. For example, an increase in K_a_ values suggests a change in some of the amino acids, and can be observed at the protein sequence level, while an increase in the K_s_ value can only be seen at the nucleotide sequence level. The K_a_/K_s_ ratio, very generally speaking, indicates the nature of the gene adaptation in reference to its predecessor; in this case, the Wuhan genes. If this ratio is greater than 1, it indicates that the gene is going through adaptational changes, or positive selection. This might imply that the gene is responding and adapting to environmental factors. On the other hand, a ratio less than 1 indicates a purifying, or stabilizing selection. In other words, it is *resisting* mutational changes, possibly indicating dependence on the amino acid sequence for function. A ratio equal to 1 indicates no change, or neutral selection. It is important to note that these interpretations are overly simplistic and are provided for the general reader who is not familiar with the concept; in reality, the situation is influenced by many factors, including biases. For example, some of those biases could be transition/transversion bias and codon-usage bias.^10^ However, for the purposes of this study, we investigated the K_a_, K_s_, and K_a_/K_s_ in genes in SARS-CoV-2 sequences collected over the course of COVID-19 as a method for identifying mutational behavior and predicting future surges.

### Daily mean values

Each of the tracked quantities (K_a_, K_s_, and K_a_/K_s_) have a number of different values associated with them calculated from the different unique sequences reported on a given day. For the ease of tracking the behavior over time, a weighted mean value was calculated for each day. Each of the accession records was assigned to a bin with a unique sequence. The calculated values were weighted by the number of sequences (records) in each of these bins. Daily average values of K_a_ and K_s_ were calculated directly. However, two methods were used for the K_a_/K_s_ ratio.

### Two alternate approaches for daily K_a_/K_s_ mean values

1. A direct approach was first used to calculate K_a_/K_s_ value for each unique sequence if the K_s_ value was not equal to 0 (the ratio K_a_/K_s_ is undefined when K_s_ is 0). However, it was found that in many cases near-zero values of K_s_ led to artificially inflated K_a_/K_s_ values that failed to intuitively communicate mutation trends. The daily weighted mean values of K_a_/K_s_ were abnormally weighted by these sequences with very small K_s_ values, resulting in very noisy behavior as a function of time. Furthermore, the ongoing curves corresponding to the K_a_/K_s_ ratio did not correspond to mean K_a_ and mean K_s_ values. It should be noted that this is not a discrepancy, but rather a consequence of the mathematically necessary exclusion of the sequences with a K_s_ value of 0. The results from this approach are in the supporting information and the values are denoted as K_a_/K_s_’.
2. As an alternative approach, we computed a daily value of K_a_/K_s_ by dividing the daily weighted mean value of K_a_ by the daily weighted mean value of K_s_. These values provide more easily interpretable results and are shown in the main manuscript.

### Forecasting future surges

The genomic surveillance data, in combination with the number of infection cases, was used to develop a monitoring approach and *a priori* forecasting of an increase in number of infections (or “surges”). K_a_, K_s_, and K_a_/K_s_ profiles were compared to the infection profile. As we previously described, the ratio of non-synonymous to synonymous mutations (K_a_/K_s_), which is typically used, did not provide clear trends. Furthermore, the rate of mutations (a derivative of observed mutations with respect to time) also did not provide a reliable prediction signal. However, we found that collective non-synonymous mutations in key proteins of SARS-CoV-2 showed a significant increase 2-4 weeks before the rapid rise in COVID-19 cases, particularly related to the surges that occurred after the emergence of the Gamma, Delta, Omicron, and BA.5 variants.

### Forecasting future make-up of variants of concern

A method to monitor the potential variants of concern (VOC) is described in Figure 2. It is based on examining isolates with genes that have K_a_/K_s_ values larger than one over a sliding 14-day window. First, the gene information for all available variants for the past 14 days are extracted, including the nucleotide and amino acid sequences and the calculated K_a_, K_s_, and their ratio values. Next, we group the data for each gene using groups of identical K_a_, K_s_, and K_a_/K_s_ values. The largest group for each gene is selected for clustering using mmseq2.^11^ Non-redundant sequences are then used for phylogenetic analysis. The potential variants are then aligned with the previous VOCs, including the reference Wuhan strain, and the amino acid substitutions are deduced using clustal-omega.^6^

**Figure 2:**
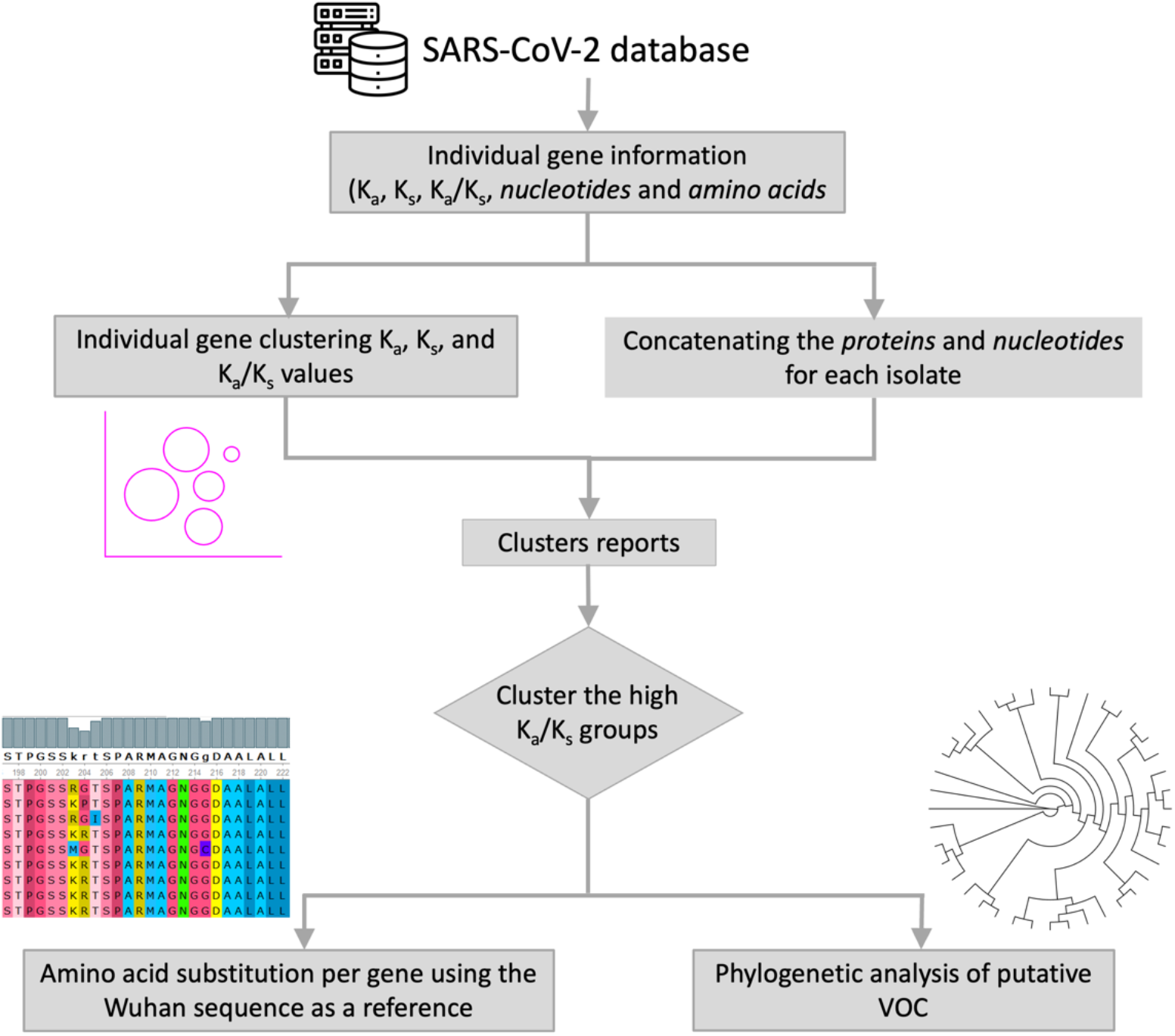
A schematic overview of the approach for predicting future mutations.

## RESULTS

### Mutational propensity of various genes

An overview of the substitutions at the nucleotide and protein level of the 26 SARS-CoV-2 proteins, since the beginning of the pandemic are depicted in Figure 3. These results are based on 8.03 million sequences sourced from GenBank ranging from the earliest days of COVID-19 to the present day. The number of quality-controlled accession records range from 2.2 million to 3.5 million for different genes (see Table S1 in the supporting information for details for each gene). The number of unique sequences as a proportion of total daily sequences ranges from almost 0% to 10.6% at the nucleotide level (Figure 3A), while at the protein level it ranges from 0.0% to 5.0% (Figure 3B). Note that the genes are listed in decreasing order of unique sequences. As expected, some of the higher numbers of unique sequences are observed for the large genes and proteins while smaller genes show some of the lowest numbers of unique sequences; however, the trends are in the middle are not uniform. Additionally, the percentages of unique proteins versus nucleotide sequences for each gene differ vastly. There is a higher percentage of unique nucleotide sequences as compared to their amino acid counterparts, indicating that most of the substitutions are synonymous.

**Figure 3:**
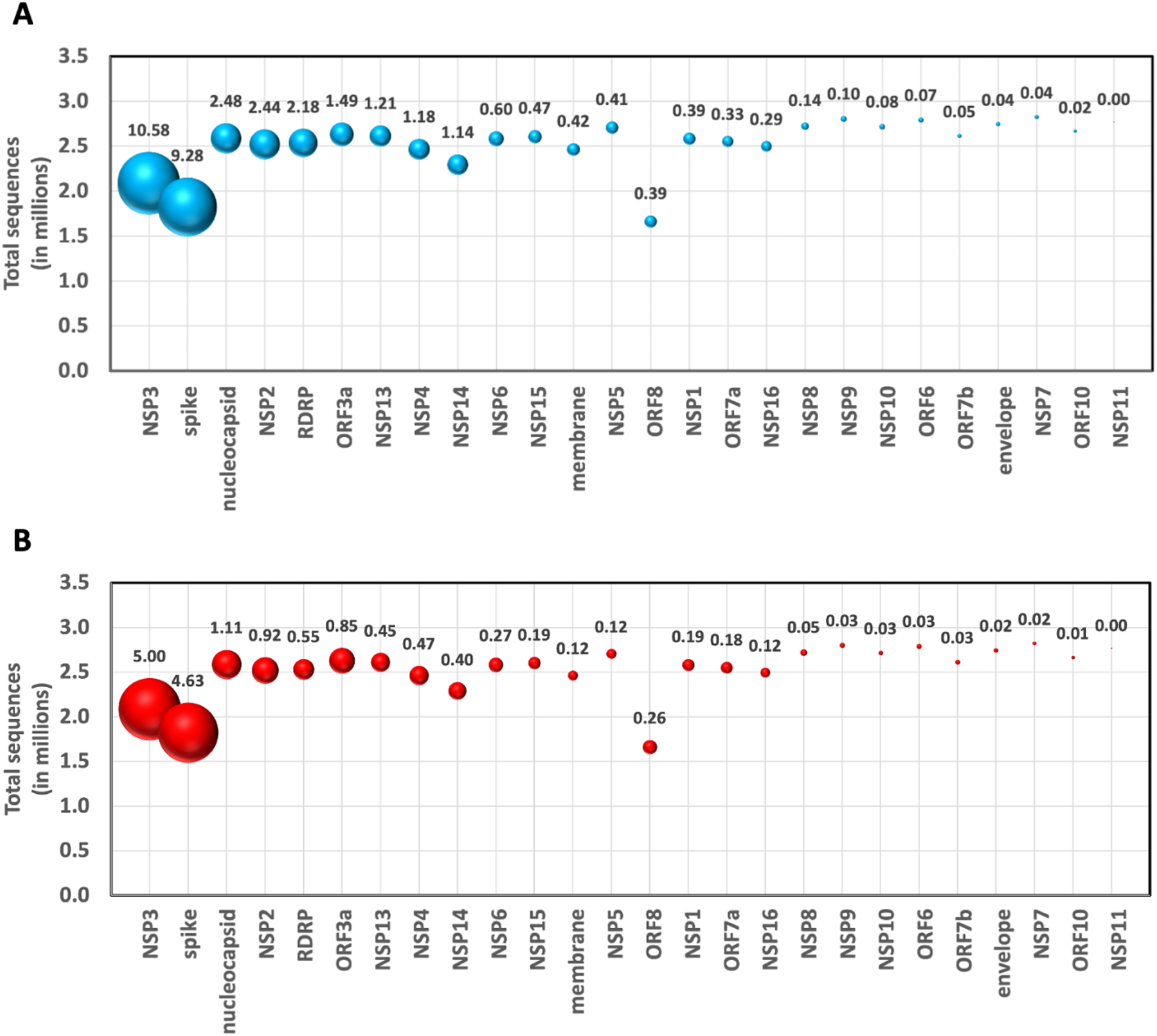
The percent of unique sequences for each gene. (A) Unique nucleotide sequences. (B) Unique protein sequences. The size of the sphere corresponds to the relative percentage of unique sequences for each gene and the percentages of unique sequences are labeled.

Mutational propensity, defined as the total number of unique mutations for each gene divided by the total length (in amino acids), displays a wide range across the genes and provides several insights (Figure 4, depicted as percentage). In the collected sequences, the spike protein, NSP6, and the nucleocapsid protein have shown the largest propensity to mutate. The spike protein is associated with viral membrane structure, while NSP6 is associated with the host’s endoplasmic reticulum membrane. Additionally, the four external proteins (spike, nucleocapsid, envelope, and membrane proteins) rank in the top six highest propensities. There are several other notable observations; most of the proteins with the lowest ranks (indicating proteins that have been more mutationally stable so far) seem to form complexes with other proteins. For example, NSP7 and NSP8 form a complex with NSP12 as part of the replication complex. Similarly, NSP10 and NSP14 are needed to activate NSP16, with NSP10 also serving as an activator of NSP14. Another notable observation is that the top-ranking proteins in mutational propensity are membrane-associated protein. This analysis can potentially be useful for protein targeting for vaccine and for drug development.

**Figure 4:**
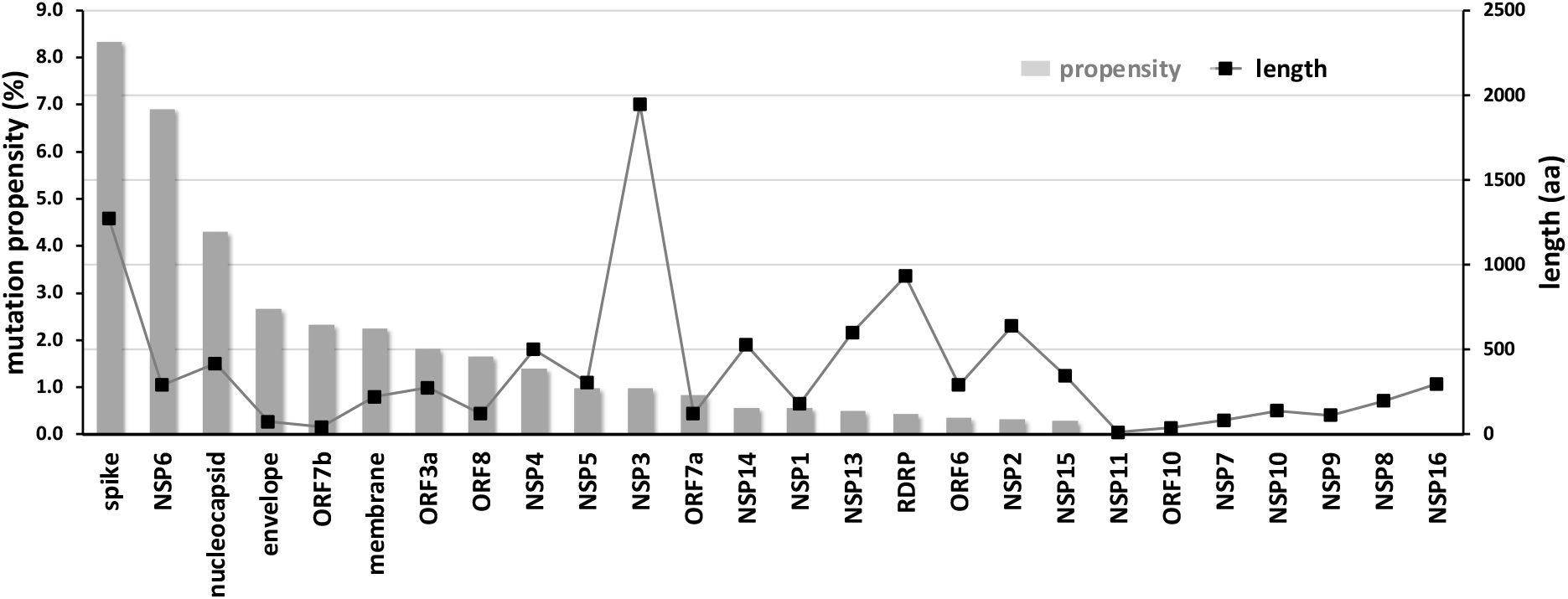
Mutational propensity of various SARS-CoV-2 genes. The number of mutations observed in each gene at the protein sequence level and normalized by sequence length is depicted (as gray bars). The genes are ranked in order of their mutational propensity. The length of each gene, shown as black lines, shows no correlation with the mutational propensity. See the table in the supporting information for the exact values.

In the remaining section, we discuss the important observations for each individual protein. We start with spike protein, as it has shown the most interesting behavior. Then we describe the three other structural proteins, followed by NSPs and the various ORFs.

### Spike Protein

This protein plays a vital role in the successful internalization of the virus into the host cells by interacting with the host cell angiotensin-converting enzyme 2 (ACE2) receptor.^12^ The protein consists of three domains: a receptor-binding domain that binds to the human ACE2, a fusion domain, and a transmembrane domain.^13^ Cleavage of the spike protein by host proteases kickstarts the process of uptake into the host cell.^14^ The spike protein has been the predominant target of mRNA vaccine development. Recently, it has been hypothesized that the spike protein may interfere with the immune synapse assembly and function that is normally responsible for coordinating the killing of virally-infected cells by cytotoxic T lymphocytes.^15^

Around 50 different mutations have been observed in the spike protein since the start of the pandemic (see Table 1 and Figure 5). Note that for our analysis, we used only the fully quality controlled sequences. As described in Table S1 in the supporting information, there are almost 2.36 million records, with a total of 227,602 unique nucleotide sequences and 115,658 unique protein sequences for this protein. Different mutations are present in all VOC but the number of mutations significantly increased beginning with the Omicron BA.1 variant in the post-vaccination period. As visible from Table 1, compared with all the other genes, the spike protein has the greatest number of observed mutations in all the SARS-CoV-2 variants. Mutations marked with an asterisk (*) are of particular interest, as these amino acids interact with ACE2 receptor, which are present in most VOCs, and again have increased since the emergence of Omicron BA.1 variant. Functionally, these mutations are expected to affect the interaction with the ACE2 receptor as a number of them are located in the receptor binding domain (Figure 5).

**Figure 5:**
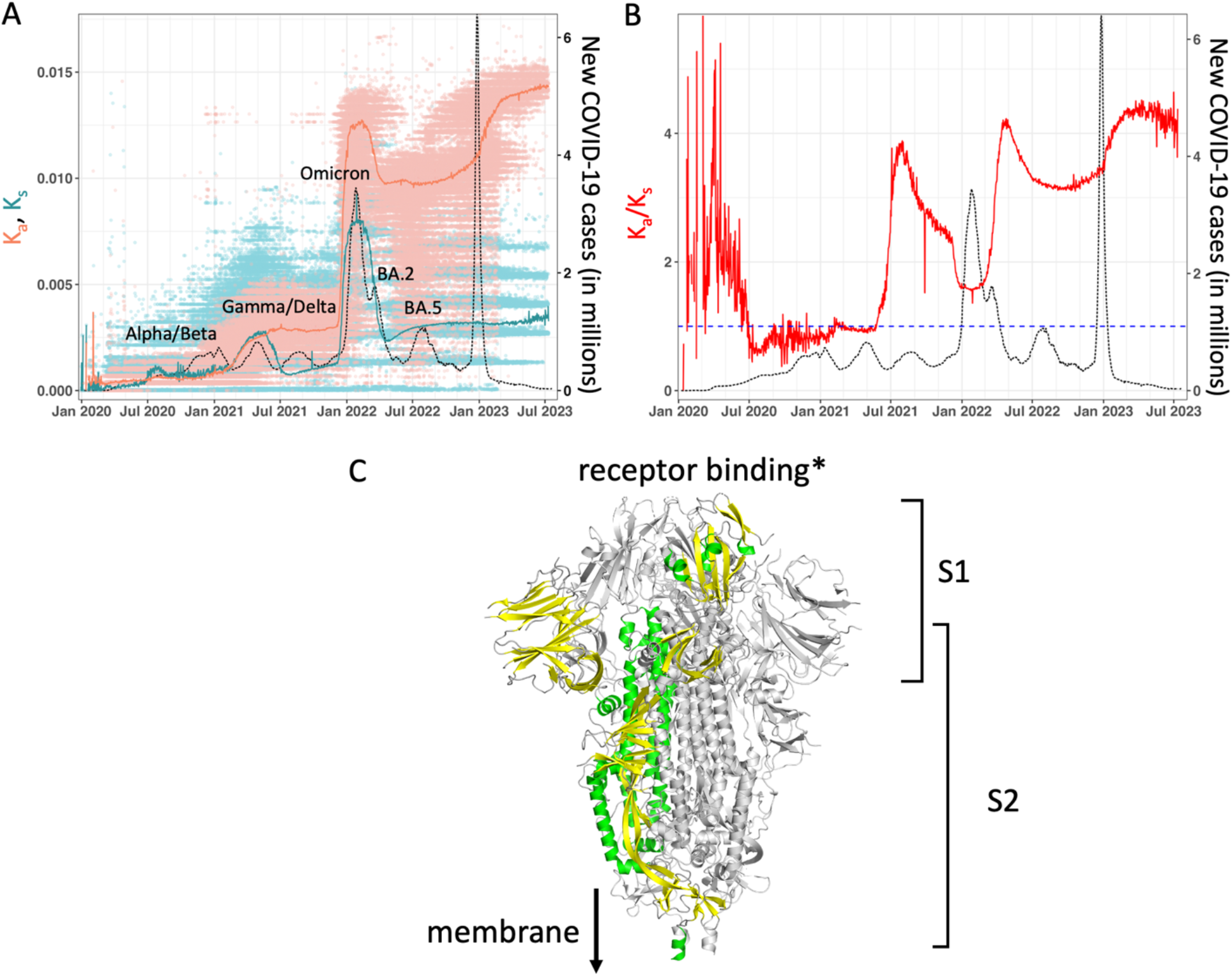
Spike protein mutation summary over the course of the COVID-19 outbreak. (A) Non-synonymous (K_a_) mutations are shown in light orange, while synonymous (K_s_) mutations are shown in cyan. Individual dots correspond to unique sequences reported on different dates. The light orange and cyan curves are mean K_a_ and K_s_ values, weighted by number of sequences observed for each unique sequence. Note that a value of 0 corresponds to a sequence identical to the Wuhan sequence, and mean weighting for each unique sequence depends upon the number of identical reported sequences that match corresponding to a particular date. The dashed black curve shows new COVID-19 cases (sliding weekly average) across the world. The peaks for COVID-19 cases are labeled with prevalent variants. Alpha/Beta, Omicron, and Omicron BA.2 and BA.5 were the prevalent variants at the time of the labeled peaks. For the two peaks in 2021, the case was less clear, with Gamma and Delta variants being observed at different times in different parts of the world. (B) Ratio of non-synonymous to synonymous mutations (K_a_/K_s_) shown in red. As in panel A, the dotted black curve shows new COVID-19 cases. (C) Structure of the spike protein, based on protein data bank (PDB) ID 6VXX.^17^ The protein is a functional trimer, with one of them shown in green and yellow color, while the other two are colored in gray. The receptor binding region (corresponds to the entries marked by * in Table 1), and subunits S1 and S2 are marked. The direction of the domain that interacts with the membrane (not shown) is marked. Note, the missing regions in the structure are indicated by dashed lines.

**Table 1:**
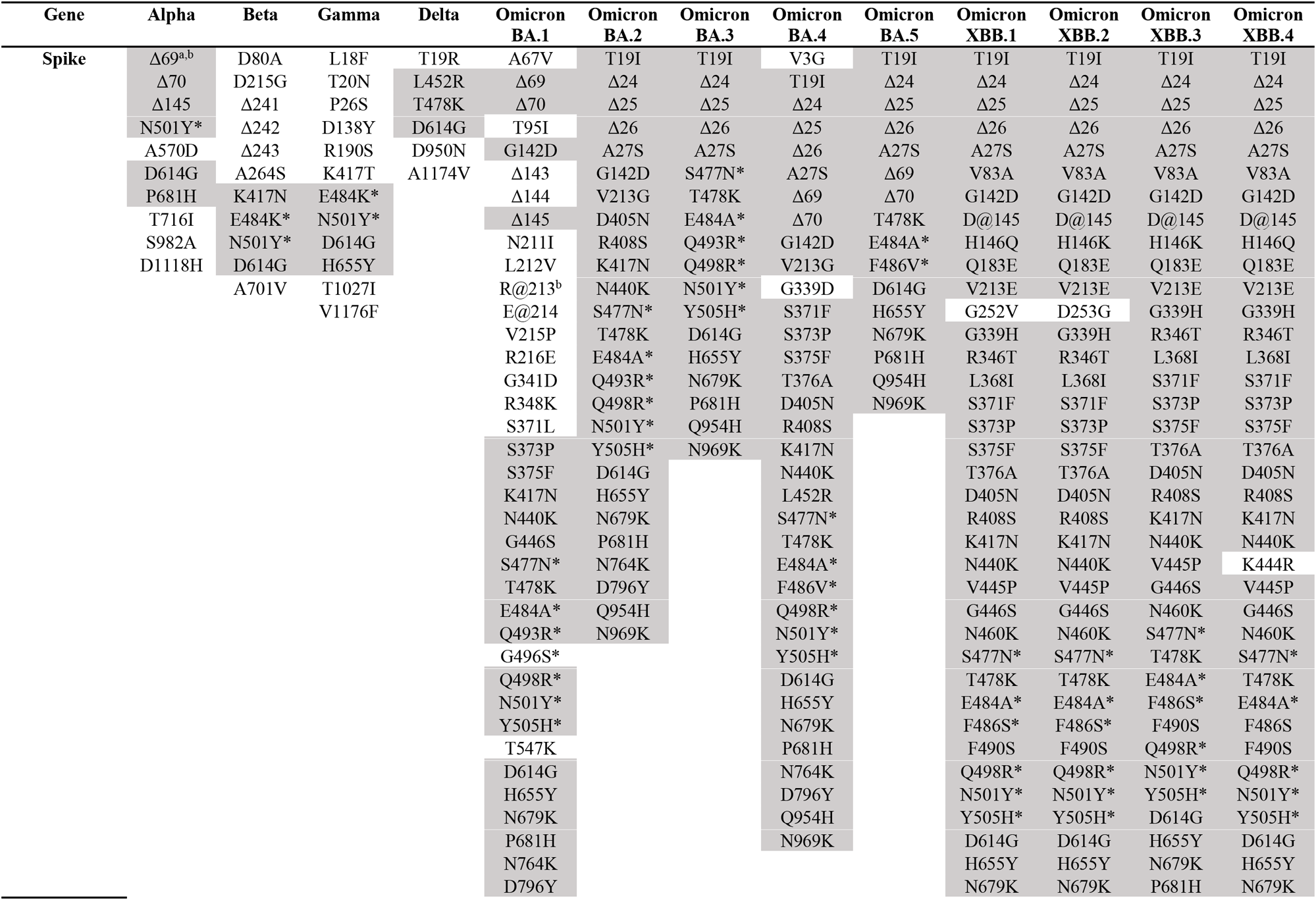
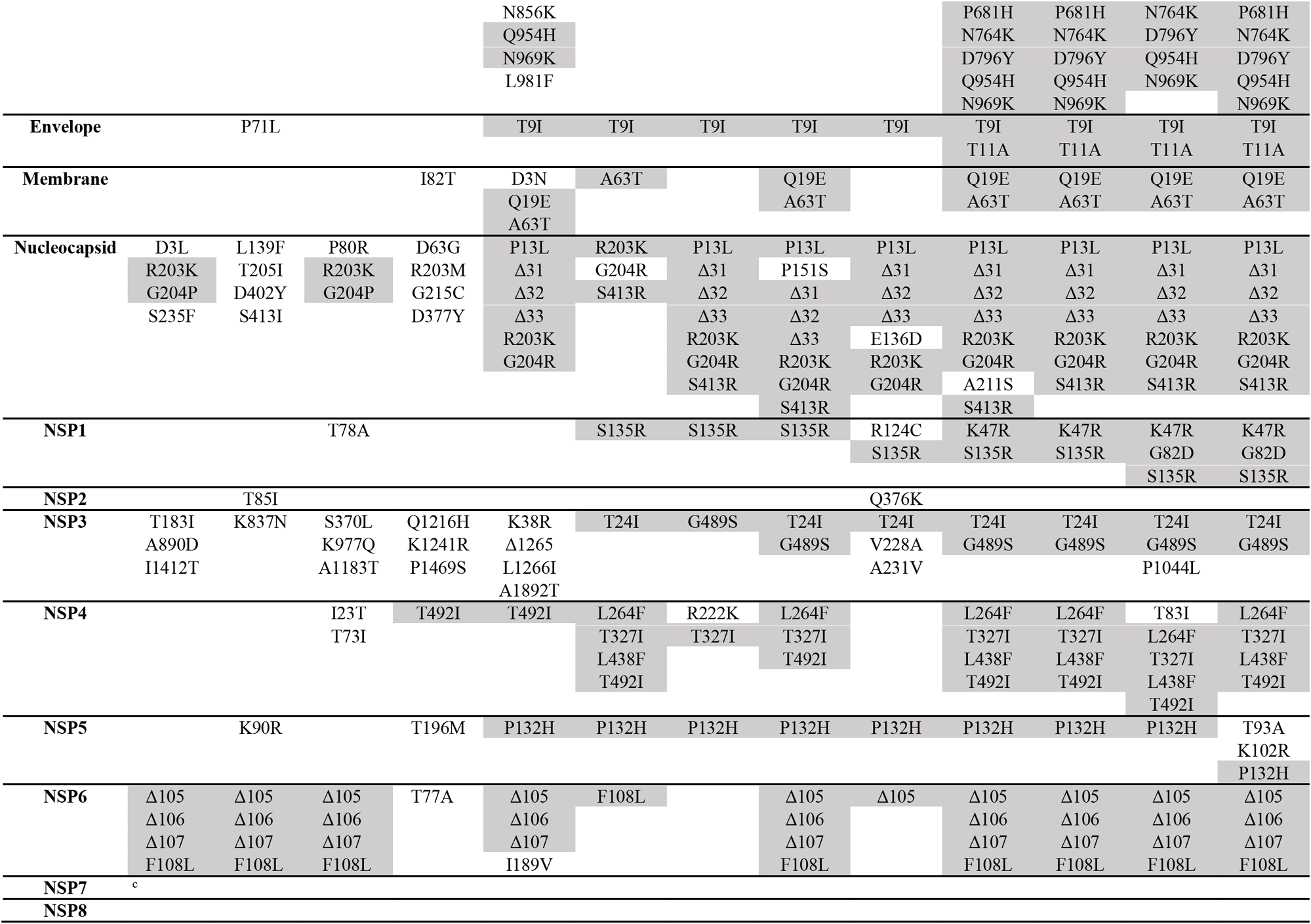
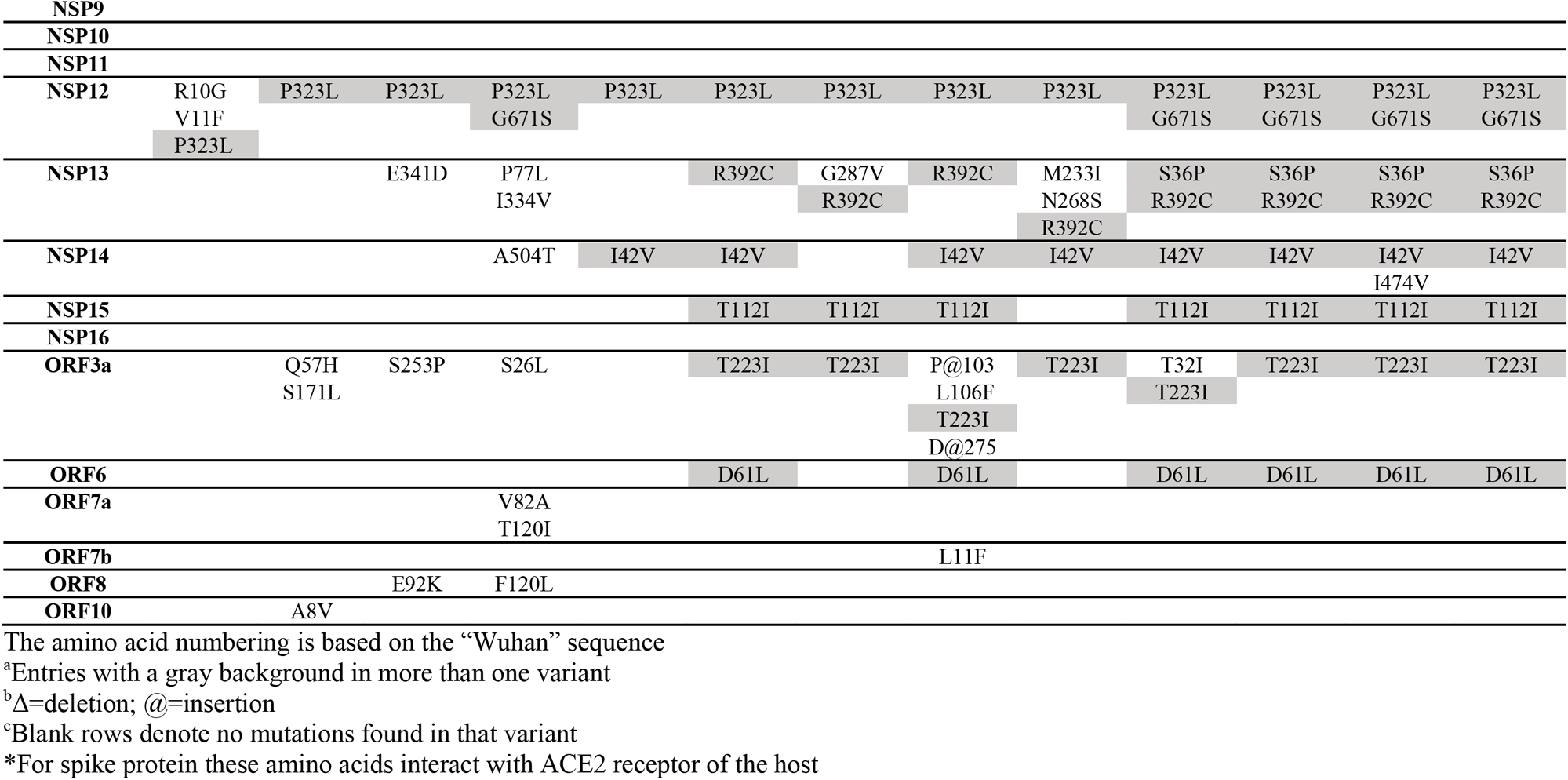
Summary of all mutations observed in the 26 SARS-CoV-2 proteins.

As a function of time the spike protein has shown a significant number of synonymous and non-synonymous mutations over the course of the COVID-19 outbreak (Figure 5A). Note that in Figure 5A, each unique value of K_a_ and K_s_ (light orange and cyan respectively) are shown as dots, which correspond to a unique sequence of the protein, while the solid orange and cyan curves are based on the weighted average values for each day. The K_a_/K_s_ ratio (red line in Figure 5B), the most commonly used metric, rose to a very high value in the initial months of the outbreak (January 2020) and has shown a tendency to decrease since then, with notable increases around July-September 2020 and May-July 2021. Note that during the initial period of the outbreak the data is substantially noisy due to the lower number of sequences, which affects the mean weighting (see Methods section for more details). The K_a_/K_s_ ratio did not show any direct relationship with the number of new COVID-19 infections (dotted black lines). However, as reported previously by our group,^16^ K_a_ showed a significant increase correlating with the Gamma/Delta variant related surge in 2021 and the Omicron surge in late 2021. A careful analysis of the data indicated that the non-synonymous mutations increased 10-14 days before each of the two surges. It should be noted that while the raw sequence data showed more unique sequences corresponding to synonymous mutations as compared to non-synonymous mutations until almost the end of 2021, the weighted non-synonymous K_a_ has showed a consistently higher value than the synonymous K_s_ since the onset of the Omicron variant in late 2021 (Figure 5A; note the difference in range between the cyan and light orange dots). The reason for this observation is that a significant number of sequences either closely resemble the Wuhan reference sequence or are weighted by more daily reports of low K_s_ values towards the beginning of the pandemic, while K_a_ is weighted more by the heavily mutated sequences that became more prevalent as time passed. Another possible explanation for this dramatic shift is that it could be a result of the adaptive pressure from the mRNA-based vaccines targeting the spike protein which were widely deployed around this time. As noticeable in Figure 5A-B, both K_a_ and K_a_/K_s_ are at their highest values as of July 2023, raising concerns about the long-term effectiveness of vaccines (and naturally acquired immunity) targeting the spike protein. The substantial number of mutations observed in Omicron XBB VOC further highlights this point. The data warrants a careful watch on the evolving situation. Note that there is large surge in COVID-19 cases between Dec. 2022 and Jan. 2023 (the largest number of COVID-19 cases reported during the course of the outbreak). The vast majority of these cases were reported from China. Information indicates that these were caused by the XBB variants, however, genomic information is not available in GenBank for these cases at the corresponding scale. Nonetheless, rise in K_a_ also corresponds to this surge in Figure 5A-B.

### Envelope protein

This protein is essential for the production and release of mature viral particles.^18^ This viroporin protein is produced in abundance throughout the course of infection, where it is thought to be most important during assembly, with a much smaller number of copies incorporated into mature virus particles.^19^ It is also believed to bind toll-like receptors of the host cell, causing it to play a key role in the hyperinflammation present in COVID-19 cases.^20^ More recent research suggests that ectopic expression of the envelope results in translational inhibition, as the viral protein binds to the initiation factor eIF2α.^21^

Amino acid substitutions for this protein are observed in the Beta variant and all the Omicron variants, which share the same mutation (Table 1). The mutation P71L, which is unique to the Beta variant, was found to have a slightly stabilizing effect on the protein structure, although its functionality is unknown^22^ (Figure 6). The T9I mutation, which is shared across all Omicron variants, dramatically reduces the selectivity of the ion channel.^23^ Due to this, when compared to the Wuhan type it will be less likely to kill the host cells and produce cytokine, which results in much less severe symptoms than variants lacking this mutation. The K_a_/K_s_ plot is extremely noisy prior to the Omicron variant surges in late December 2021 (Figure 6B), and it shows a steep increase during this time and remains elevated to present day. This is also reflected in the K_a_ plot, which showed the most significant increase around the Omicron surge and further started increasing recently.

**Figure 6:**
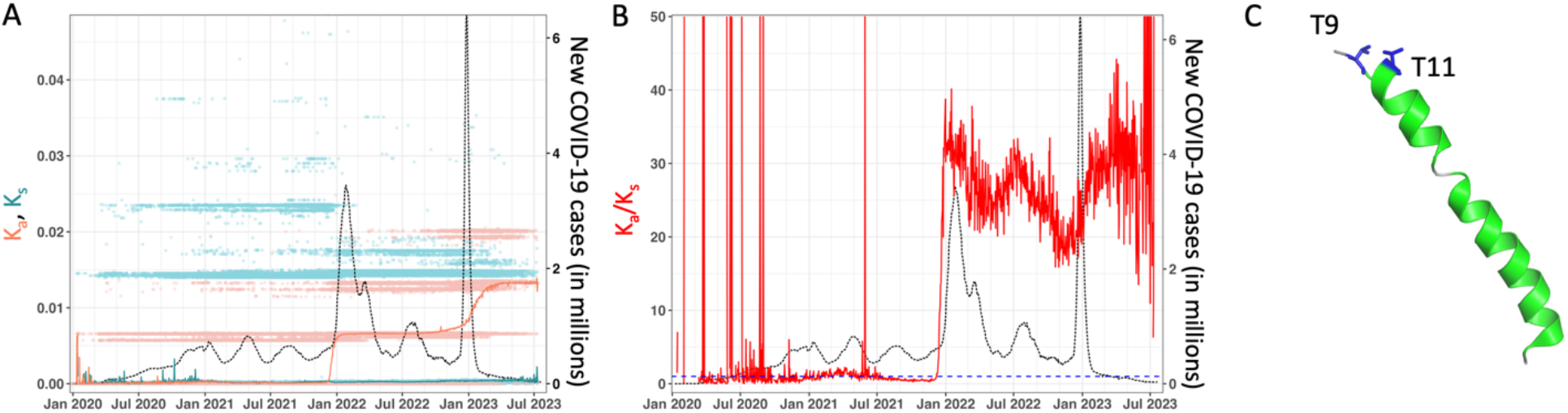
Envelope protein mutation summary over the course of the COVID-19 outbreak. (A) Non-synonymous (K_a_) and synonymous (K_s_) mutations. See Figure 5 for the VOC names corresponding to the peaks in infections and other details. (B) Ratio of non-synonymous to synonymous mutations (K_a_/K_s_). (C) Structure of the envelope protein, based on protein data bank (PDB) ID 7K3G.^24^ The two residues T9 and T11 are shown, which show mutations since the Omicron VOCs. Note the residue 71 that showed mutation in Beta variant is located in the region where the situation information is not available.

### Membrane protein

The membrane protein is believed to interact with the nucleocapsid protein in the formation of mature viral particles.^25^ It is the most abundant protein in virions and has been found to be involved in a host of different functions, including but not limited to interfering with the host interferon system, inducing autophagy, serving as a protective antigen, and serving as a viroporin.^19^

Interestingly, for the membrane protein Q19E and A63T amino acid substitutions are observed in most of the Omicron subvariants, while the other variants, with the exception of the Delta variant, did not accumulate any mutations (Figure 7 and Table 1). The I82T mutant, which appeared in the Delta variant, has a slight stabilizing effect on the protein structure.^26^ It is assumed that his mutation enhances glucose uptake during viral replication.^27^ The K_a_ values show that amino acid substitutions started at the Delta variant surge in June 2021 and have remained high since then (Figure 7A). While the Beta variant’s sequence shows a single amino acid substitution, it is not obvious on the K_a_ plot, likely due to how noisy the plot is at that region. The K_a_ and K_a_/K_s_ plots show rise 2-4 weeks before the increase in number of infections for several VOCs (see Figure 7A-B). However, compared to other proteins, the K_a_/K_s_ has shown a significant decrease recently.

**Figure 7:**
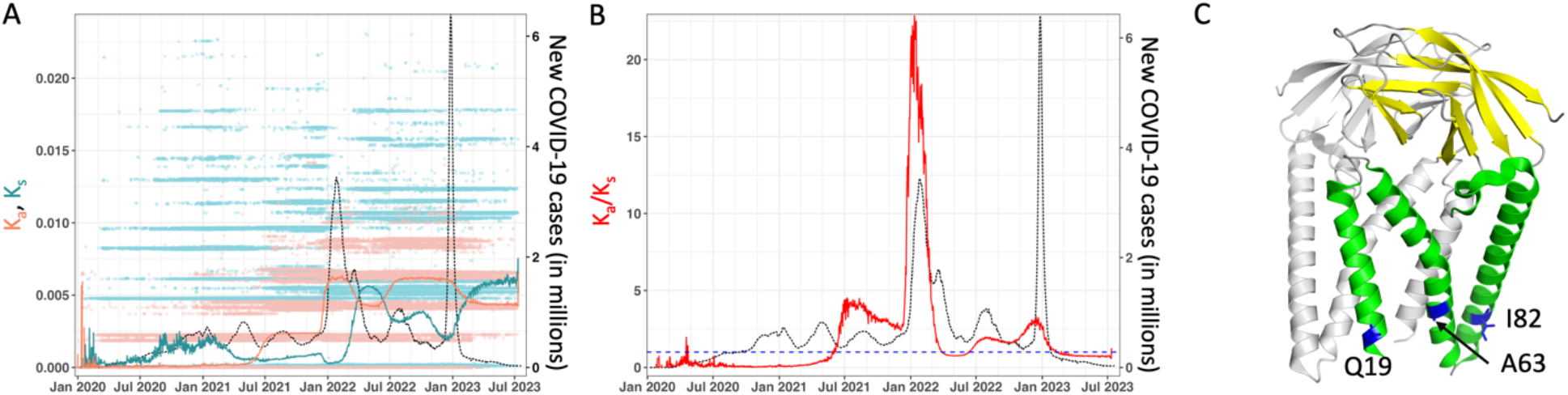
Membrane protein mutation summary over the course of the COVID-19 outbreak. (A) Non-synonymous (K_a_) and synonymous (K_s_) mutations. See Figure 5 for the VOC names corresponding to the peaks in infections and other details. (B) Ratio of non-synonymous to synonymous mutations (K_a_/K_s_). (C) Structure of the membrane protein, based on protein data bank (PDB) ID 8CTK.^28^ The structure is shown as a dimer, with only one protomer colored in green and yellow. The location of three important residues showing mutations are marked.

### Nucleocapsid

This protein is essential for packaging the viral genome into virion particles.^29^ Specifically, this protein serves as the physical link between the positive-RNA genome and the envelope.^30^ It has two domains; the N-terminal domain that binds to the viral RNA and the C-terminal domain that binds to the M-protein in the envelope with a large disordered region in the middle.^31^ The other aspect of the nucleocapsid protein’s function, an unknown, is its interaction with NSP3, suggesting a role in the viral life cycle.^31^ A recent study has also correlated amino acid substitutions in this protein to the immune response and concluded that these mutations are correlated to the immune system evasion attempts, suggesting these sites as drug targets.^32^

Several mutations in this protein have been observed in all the variants (Figure 8 and Table 1). The double mutant R203K/G204R is present in all Omicron variants, with the Alpha and Gamma variants containing a similar double mutant, R203K/G204P. It was shown that this mutation augments the nucleocapsid phosphorylation, which in turn increases replication.^33^ Interestingly, this mutation also increases the resistance to inhibition of the GSK-3 kinase. The Delta variant mutation G215C is located near the normally disordered region of the protein-protein interface (Figure 8C). Once introduced, this mutation causes the stabilization of the transient helix, enhancing the inter-protein interaction, which leads to the formation of a tetrameric rather than dimeric state. The nucleocapsid tetramer has a higher binding affinity to nucleic acid, which increases the virus’ efficiency.^34^

**Figure 8:**
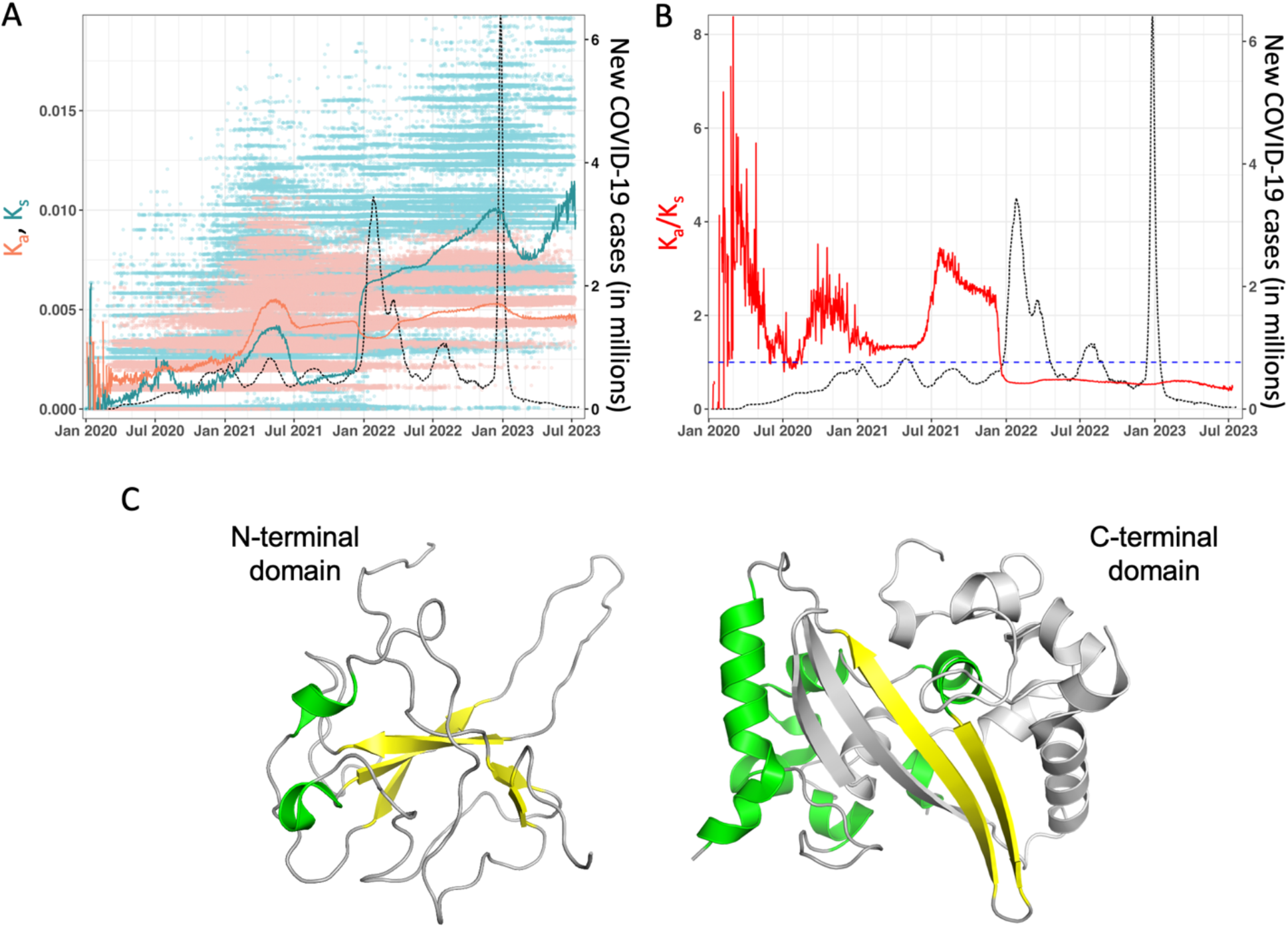
Nucleocapsid protein mutation summary over the course of the COVID-19 outbreak. (A) Non-synonymous (K_a_) and synonymous (K_s_) mutations. See Figure 5 for the VOC names corresponding to the peaks in infections and other details. (B) Ratio of non-synonymous to synonymous mutations (K_a_/K_s_). (C) Structure of the nucleocapsid protein, based on protein data bank (PDB) IDs 6YI3^35^for N-terminal domain and 7O05^36^ for the C-terminal domain. The linker region between the two domains is not shown. The C-terminal domain structure is shown as a dimer, with only one protomer colored in green and yellow.

From the broader genomic surveillance perspective, aside from the spike protein, the nucleocapsid protein demonstrates one of the highest propensities for mutation among the SARS-CoV-2 genes. Earlier in the pandemic, during the Alpha, Beta, Gamma, and Delta variant surges, the K_a_/K_s_ ratio indicates that the protein was under positive selection, indicating some non-conservative amino acid substitutions (Figure 8B). However, starting at the Omicron subvariant surges, with the exception of the BA.2 subvariant, the K_a_/K_s_ ratio switched to indicate a purifying pressure compared to the Wuhan strain. The K_a_ and K_s_ plots demonstrate the progression of the values though the surges. The ratio value remained larger than 1 until the Omicron BA.1 variant, when it dropped and remained low since.

### NSP1

Literature reports suggest that NSP1 is involved in repressing host transcription and preventing interferon induction.^37^ Recent structural work has proposed its mechanism of action involves the last 33 amino acids of the C-terminal domain of the protein tightly binding to the 40S ribosomal subunit, physically blocking mRNA from entering the channel and therefore preventing translation.^38–40^ It has further been hypothesized that NSP1 skews kinetics in its favor by uncapping cellular mRNA, leading to its degradation, while also recognizing a viral 5’ UTR stem loop to override the translation block.^41^ Moreover, NSP1, along with NSP14, has been shown to inhibit the accumulation of the human long interspersed element 1 (LINE-1) open reading frame ORF1p, suggesting a direct interfering with the interferon response.^42^

Figure 9 shows the summary of mutation behavior for NSP1 over the course of the COVID-19 outbreak. The K_a_ and K_s_ values show typical trends of increases over time. However, no correlation between mutation values and increase in number of new COVID-19 infections is particularly observable for NSP1. K_s_ did show a slight increase around April 2021; however, no significant increase was observed in K_a_ values in the initial period of the pandemic. This signals nucleotide-level changes in NSP1 sequences, with silent mutations that cause no changes in the amino acid sequence at the protein level. K_a_ and the K_a_/K_s_ ratio did show a significant increase after the Omicron BA.1 surge (January 2022 onwards). This increase coincides with the most common S135R amino acid substitution observed in all Omicron variants except BA.1 (Table 1). Not much is known about the functional importance of this and other mutations. Structural information is only available from residues 13 to 127 (Figure 9C), with observed mutations shown. Note that S135R falls outside the known structure.

**Figure 9:**
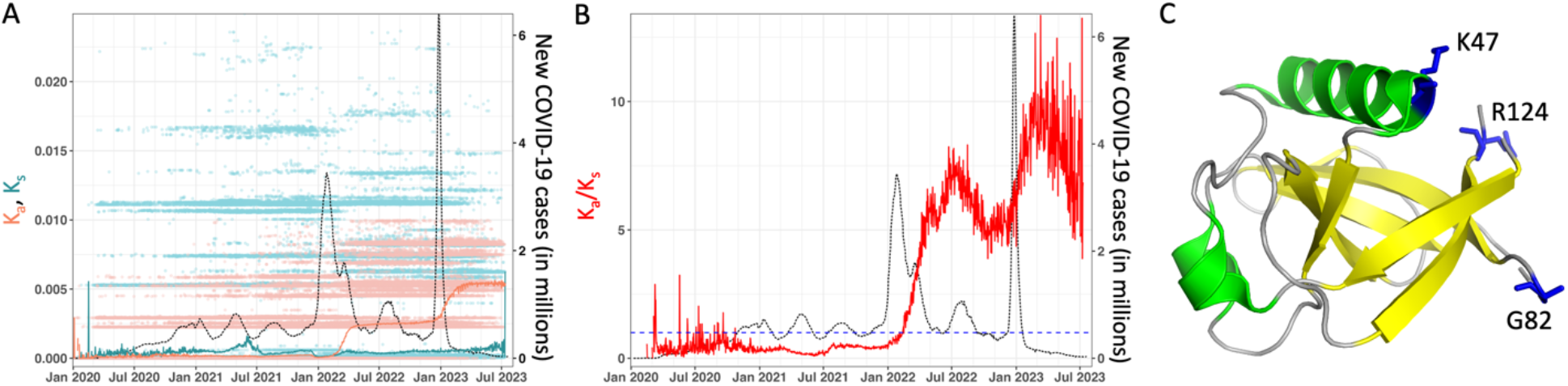
NSP1 mutation summary over the course of the COVID-19 outbreak. (A) Non-synonymous (K_a_) and synonymous (K_s_) mutations. See Figure 5 for the VOC names corresponding to the peaks in infections and other details. (B) Ratio of non-synonymous to synonymous mutations (K_a_/K_s_). (C) Structure of NSP1, based on protein data bank (PDB) ID 7K3N.^43^ Location of residues showing mutations are marked, note S135 is not listed as it falls outside the region 13-127 for which structural information is available.

### NSP2

Research suggests this protein might be involved in viral genome replication.^44^ This is supported by a study finding NSP2 to be found in the perinuclear foci and co-localized with the nucleocapsid protein, suggesting a presence at the site of viral RNA replication.^45^ More recently, a function involving the inhibition of miRNA silencing pathways has been suggested, allowing for NSP2 to suppress the host immune system through taking control of posttranscriptional silencing.^46^

Examination of the amino acid alignments of the major variants showed single, unique mutations in the Beta and Omicron BA.5 variants (Figure 10 and Table 1). One such mutation, T85I, is located on the protein’s surface (Figure 10C). The substitution of a hydrophobic residue in the place of a polar residue indicates a potential for destabilization of the structure. Indeed, in accordance with a computational study, the T85I mutant is more flexible and less stable than the Wuhan-type protein.^47^ Another mutation observed in Omicron BA.5 was Q376K, however, this mutation was not observed in other VOCs. Despite the noisy nature of the K_a_/K_s_ plot, the K_a_/K_s_ ratio decreases sharply after January 2021, around the end of the beta surge (Figure 10B).

**Figure 10:**
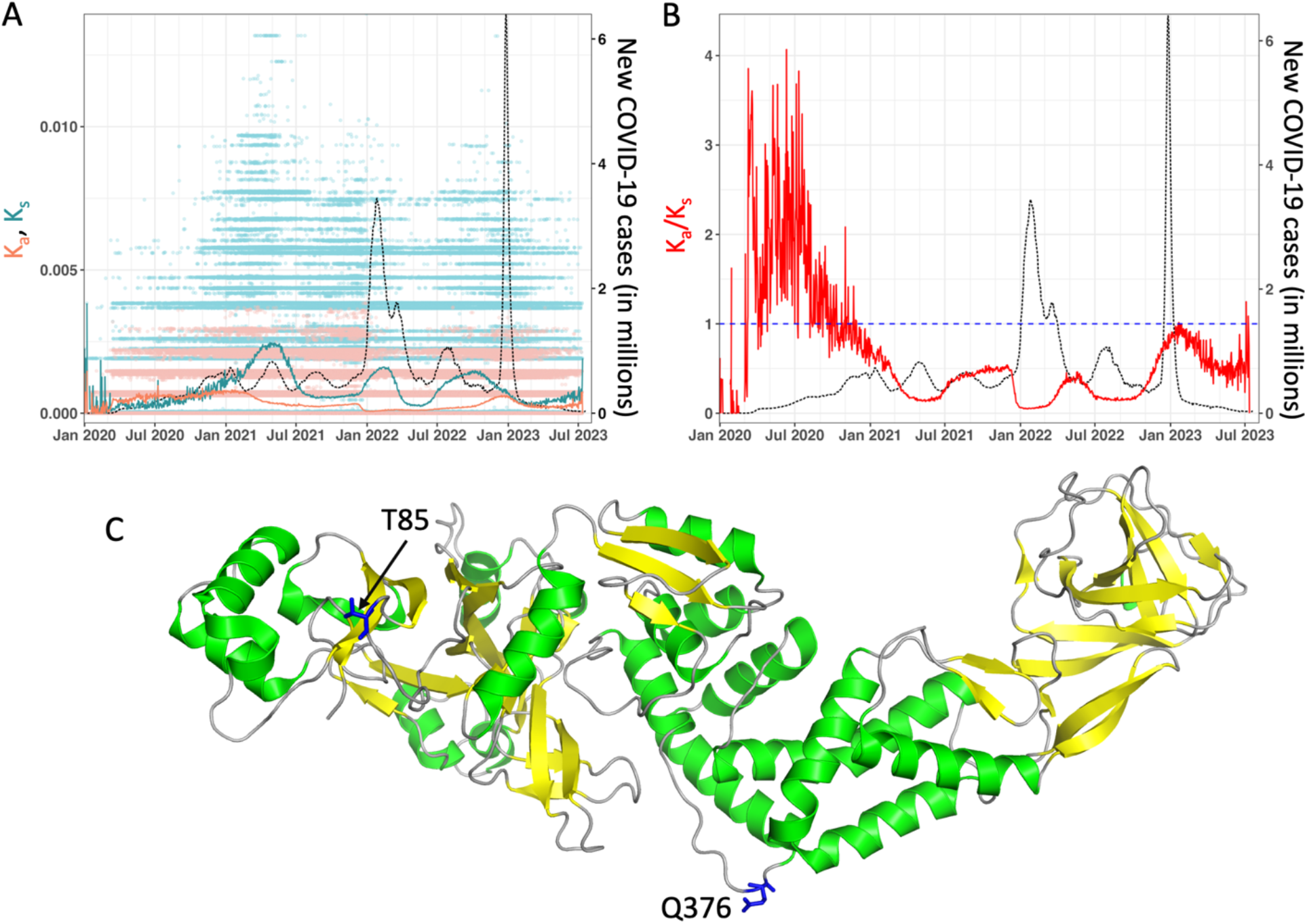
NSP2 mutation summary over the course of the COVID-19 outbreak. (A) Non-synonymous (K_a_) and synonymous (K_s_) mutations. See Figure 5 for the VOC names corresponding to the peaks in infections and other details. (B) Ratio of non-synonymous to synonymous mutations (K_a_/K_s_). (C) Structure of NSP2, based on protein data bank (PDB) ID 7MSW [DOI:10.2210/pdb7MSW/pdb].

### NSP3

This is the largest protein in SARS-CoV-2 with multiple domains (see Figure 11). Research has shown that it interacts with the nucleocapsid protein.^31^ However, the function of this interaction is still not fully understood. It is also speculated that this protein, along with NSP5 and NSP14, might be involved with maintaining an optimal environment for viral RNA synthesis, as research demonstrated their ability to repress the expression of the reverse transcriptase activity of the LINE-1 open reading frame ORF2p.^42^ Additionally, NSP3 pairs with NSP4 in the formation of replication organelles for the virus.^48^ Kahn et al have shown that residues 499 to 533 on NSP3 interact with the nucleocapsid protein, making them a potential target for drugs.^31^ Supporting this potential usage, none of those residues seem to be substituted in any of the variants, suggesting that they are essential for its function (Table 1). Additionally, NSP3 is a papain-like cysteine protease (PLPro) that processes the viral polyprotein.^49^ There is active research to develop inhibitors for this protein; specifically, targeting its conserved macro domain (Mac1), as it is critical for pathogenesis of SARS-CoV-2.^50^

**Figure 11:**
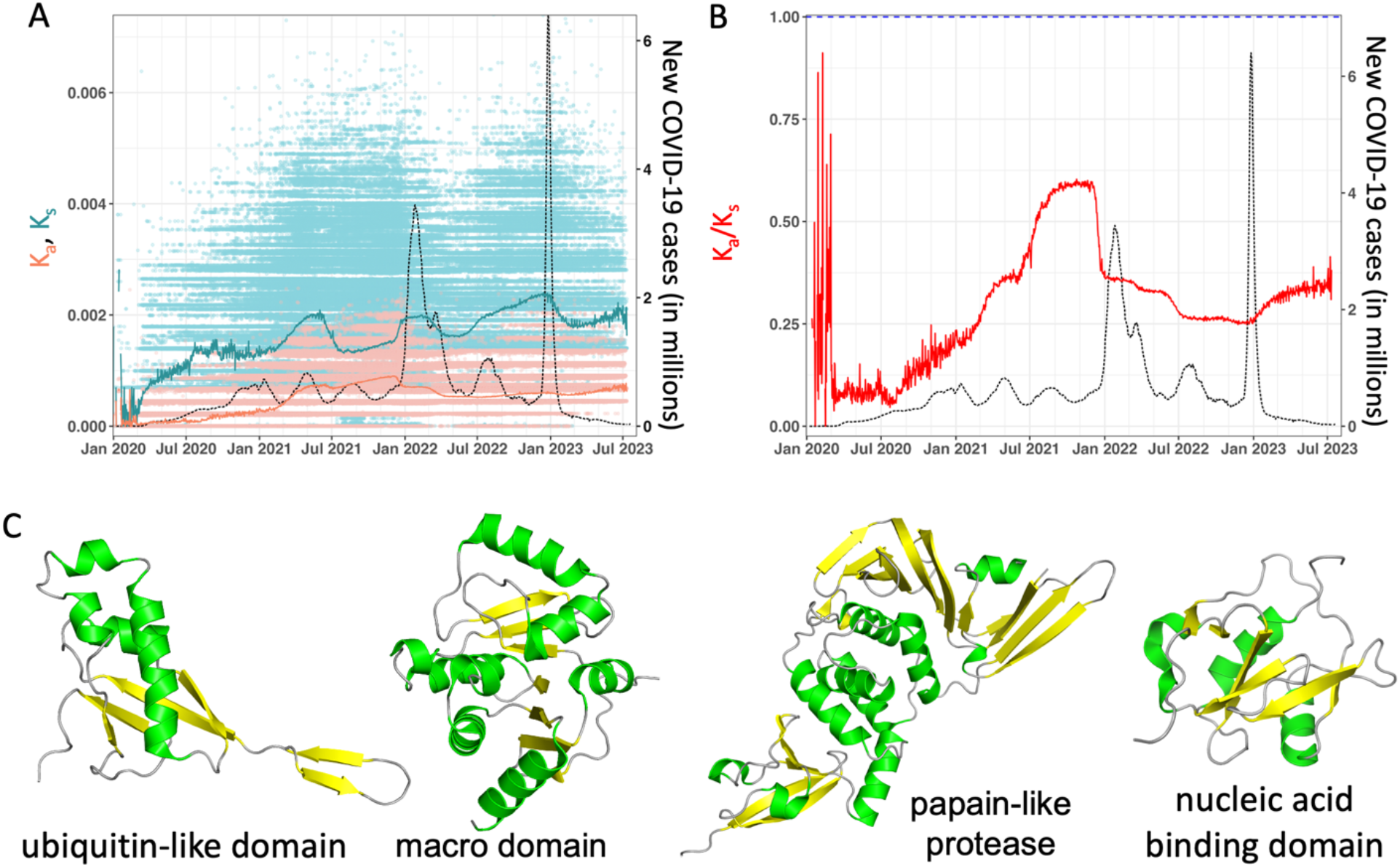
NSP3 mutation summary over the course of the COVID-19 outbreak. (A) Non-synonymous (K_a_) and synonymous (K_s_) mutations. See Figure 5 for the VOC names corresponding to the peaks in infections and other details. (B) Ratio of non-synonymous to synonymous mutations (K_a_/K_s_). (C) Structure of NSP3, based on protein data bank (PDB) 7KAG [DOI:10.2210/pdb7KAG/pdb] (ubiquitin-like domain, Ubl1), 6YWK (macro domain, Mac1),^52^ IDs 7CMD (papain-like protease, PLpro),^51^ and 7LGO [DOI:10.2210/pdb7LGO/pdb] (nucleic acid binding domain).

Amino acid substitutions lack a general pattern observed throughout all the VOCs, with many variants containing unique mutations. The substitutions T24I and G489S are the only mutations to occur more than once, with concurrent occurrence in all variants except Omicron BA.5 beginning with Omicron BA.4. The K_a_ data is reflective of the adaptational mutations we observed (Figure 11A). At the start of March 2021, the average K_a_ values of the isolates were higher for the Gamma and Delta variants. The values started to decrease in December of 2021, but still remained higher than the background.

### NSP4

This gene encodes an endoplasmic reticulum-bound protein.^54^ Comparative studies suggest that this protein may play a role in the virus replication assembly process.^55^ Specifically, NSP4 has been linked to intracellular double-membrane vesicle formation, with NSP4 working in conjunction with NSP3 to perform membrane pairing.^56^ More recently, it has been shown that the endoplasmic reticulum proteins VMP1 and TMEM41B are essential, as they facilitate NSP3 and NSP4, in conjunction with the ER, to generate the replication organelles.^48^

No substitutions were observed in the Alpha, Beta, and Omicron BA.5 variants, but amino acid substitutions were observed in all other VOCs (Table 1). Interestingly, the Delta and Omicron BA.1, BA.2, and BA.4 variants all share the T492I substitution, making it the most common substitution. This residue is located in the M-domain, within the lipid bilayer. Note, structural information is not available for NSP4 from SARS-CoV-2. Due to the increase in hydrophobicity caused by this mutation, it is suggested that this mutation stabilizes the RTC, enhancing replication.^57^ The T327I mutation, found in Omicron variants BA.2-BA.4, is suspected to function similarly. The K_a_/K_s_ plot suggests an increase in non-synonymous mutations around January 2022 (Figure 12B). This plot, in conjunction with the separate K_a_ and K_s_ plots, suggest that the number of nonsynonymous substitutions are increasing, starting around the Omicron BA.1 surge. The observed amino acid substitutions echo the plots’ trajectories. It seems that the Omicron subvariants mark the beginning of retaining the same core set of substitutions.

**Figure 12:**
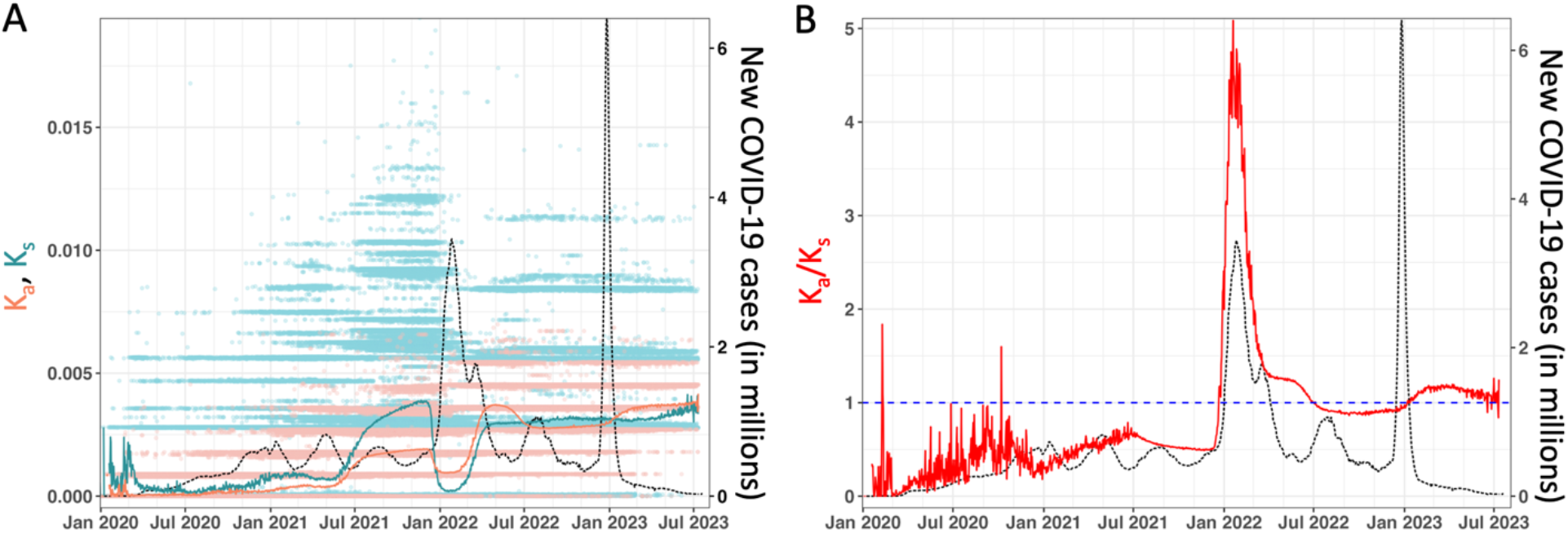
NSP4 mutation summary over the course of the COVID-19 outbreak. (A) Non-synonymous (K_a_) and synonymous (K_s_) mutations. See Figure 5 for the VOC names corresponding to the peaks in infections and other details. (B) Ratio of non-synonymous to synonymous mutations (K_a_/K_s_). As in panel A, the red curve shows new COVID-19 cases.

### NSP5

This protein is a 3C-like proteinase. It processes the polyproteins PP1a and PP1ab into their individual constituent proteins.^58^ It also is believed to play a role in evading the host’s immune system, as it also cleaves NF-kB essential modulator (NEMO) at multiple sites, ultimately resulting in inhibition of the production of interferon-beta (IFN-b).^59^ Recent work suggests that NSP5 also works with NSP3 and NSP14 to repress the expression of the reverse transcriptase activity of the LINE-1 open reading frame ORF2p.^42^

Five unique amino acid substitutions were observed in this protein (Figure 13 and Table 1): K90R, which is observed only in the Beta variant; T196M, which is observed only in the Delta variant; T93A and K102R, which are observed only in the Omicron XBB.4 variant; and P132H, which is observed in all Omicron variants. No amino acid substitutions were observed in the Alpha or Gamma variants. The K90R mutation, located in domain I of the protein, has shown functional relevance despite both residues containing polar, positive sidechains (Figure 13C). In one study, the catalytic efficiency of this mutant is 56% that of the Wuhan type.^60^ The P132H mutation, located in domain II, displays a different effect. Despite a 2.6°C lower thermal stability,^61^ the enzymatic activity is 44% higher than that of the Wuhan strain.^60^

**Figure 13:**
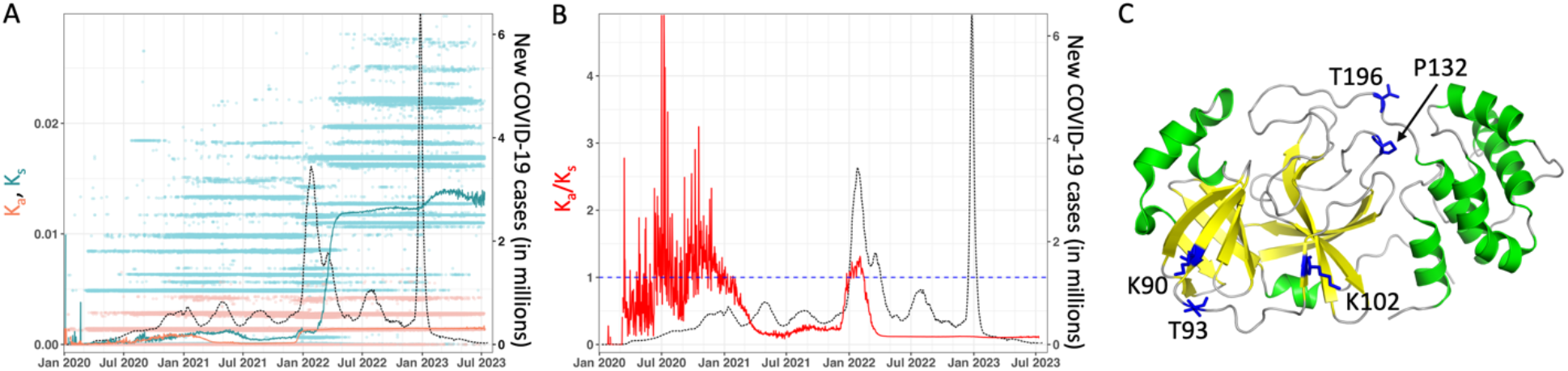
NSP5 mutation summary over the course of the COVID-19 outbreak. (A) Non-synonymous (K_a_) and synonymous (K_s_) mutations. See Figure 5 for the VOC names corresponding to the peaks in infections and other details. (B) Ratio of non-synonymous to synonymous mutations (K_a_/K_s_). (C) Structure of NSP5, based on protein data bank (PDB) ID 7LYH.^62^

The plots, and particularly the data for K_a_, demonstrate two distinct regions of increase in the nonsynonymous amino acids’ substitutions (Figure 13). The first region spans from the start of the pandemic until February 2021; this region includes the both the Alpha and Beta variant surges. The second region begins around mid-December 2021 and runs until present day. Analyzing the amino acid substitutions confirm that all the Omicron variants have both higher nonsynonymous and synonymous substitutions. The nonsynonymous mutations can be seen clearly at the nucleotide sequence level.

### NSP6

This protein is an endoplasmic reticulum membrane protein that regulates autophagy of the viral particles, which ultimately allows the viral particles to be successfully exported outside the host cell.^63^ Hence, NSP6 is an essential part of protecting viral particles. In corroboration with this, a study correlated a single nucleotide polymorphism, the substitution L37F, to destabilization of an NSP6 fold that results in decreased virulence.^64^ Indeed, recent research suggests that NSP6 and the spike proteins are essential for the Omicron subvariants’ attenuation, which further suggests its role in evasion of the immune system.^65^ Remarkably, it has been shown that NSP6 carries different sets of mutations based on geographical locations.^66^

Interestingly, the studied L37F substitution was not observed in any of the variants of concern; however, other changes were observed (Table 1). Three-amino acid-long deletions at positions 105, 106, and 107 were observed in all variants except for the Delta and Omicron BA.2, BA.3, and BA.5 variants. Additionally, three additional amino acid substitutions were observed in different variants, with the F108L mutation shared by several different variants. The structure of NSP6 from SARS-CoV-2 is not available. The K_a_ and K_a_/K_s_ plots suggest a high number of nonsynonymous substitutions February 2021, around the gamma surge, and December 2021, at the Omicron surge (Figure 14B). It also displays a lower K_a_/K_s_ ratio around August of 2021, coinciding with the Delta variant surge. This trend is also reflected in the dearth of amino acid substitutions in Delta variant.

**Figure 14:**
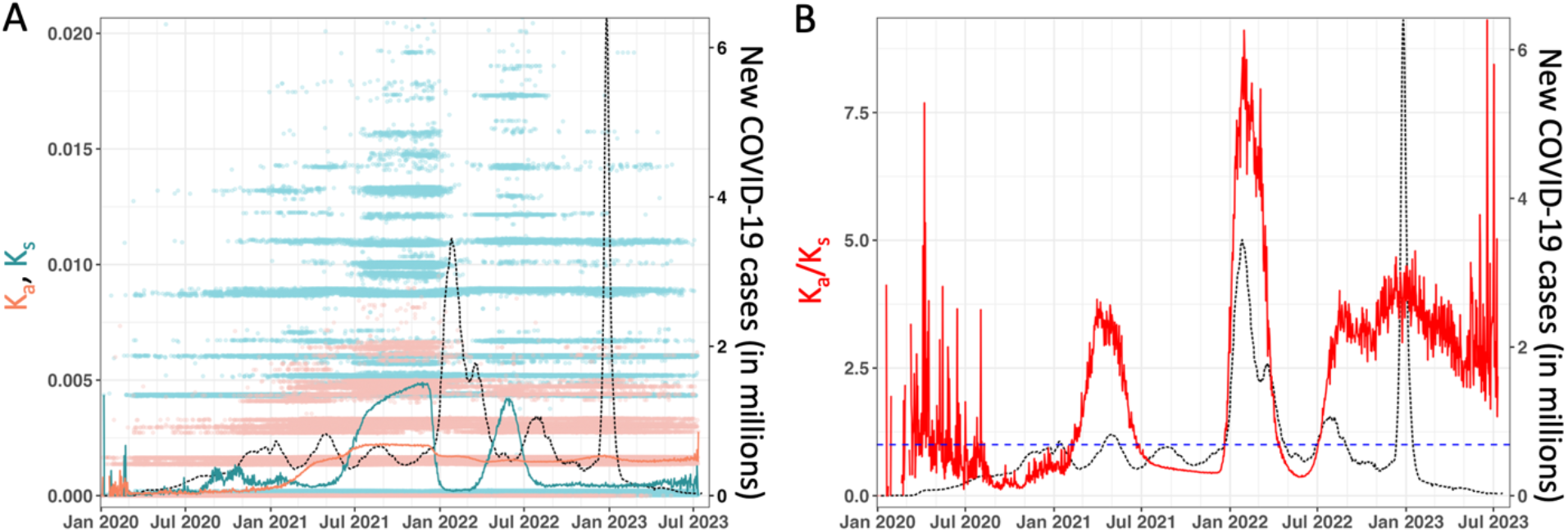
NSP6 mutation summary over the course of the COVID-19 outbreak. (A) Non-synonymous (K_a_) and synonymous (K_s_) mutations. See Figure 5 for the VOC names corresponding to the peaks in infections and other details. (B) Ratio of non-synonymous to synonymous mutations (K_a_/K_s_).

### NSP7

This protein is a cofactor of the replication complex, which includes NSP12 (RNA-dependent RNA polymerase) and NSP8 proteins.^67^ Hence, it plays an essential role in viral genome replication and transcription. It most closely interacts with NSP8, which it forms a super-complex with, that is a critical cofactor for NSP12.^68^ Interestingly, no amino acid substitutions have been observed in any of the variants (Table 1). This suggests that this gene is under little to no adaptation pressure, possibly due to its high interactivity with other viral proteins. The K_a_ and the K_s_ plots show that there are no nonsynonymous amino acid substitutions of note (Figure 15).

**Figure 15:**
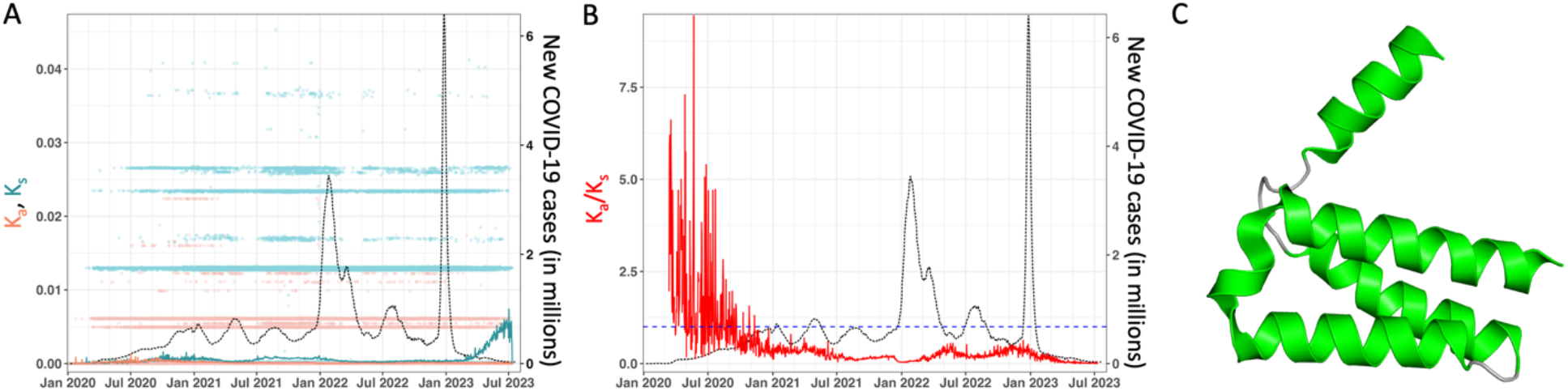
NSP7 mutation summary over the course of the COVID-19 outbreak. (A) Non-synonymous (K_a_) and synonymous (K_s_) mutations. See Figure 5 for the VOC names corresponding to the peaks in infections and other details. (B) Ratio of non-synonymous to synonymous mutations (K_a_/K_s_). (C) Structure of NSP7, based on protein data bank (PDB) ID 7JLT.^69^

### NSP8

This protein, along with NSP7, is a cofactor for the NSP12 enzyme. Studies suggest that the transcription complex is a hetero-tetramer composed of one molecule of NSP7, one molecule of NSP12, and two molecules of NSP8.^67^ Like NSP7, no amino acid substitutions were observed in any variant, suggesting that those proteins are under little to no pressure to adapt (Table 1). Additionally, the same observation of the mutation pattern is noted in the individual and K_a_/K_s_ plots for this protein (Figure 16). Of note, the K_s_ plot suggests a sharp increase in synonymous nucleotide substitutions, as it shows an increase after the Omicron BA.2 surge after April 2022.

**Figure 16:**
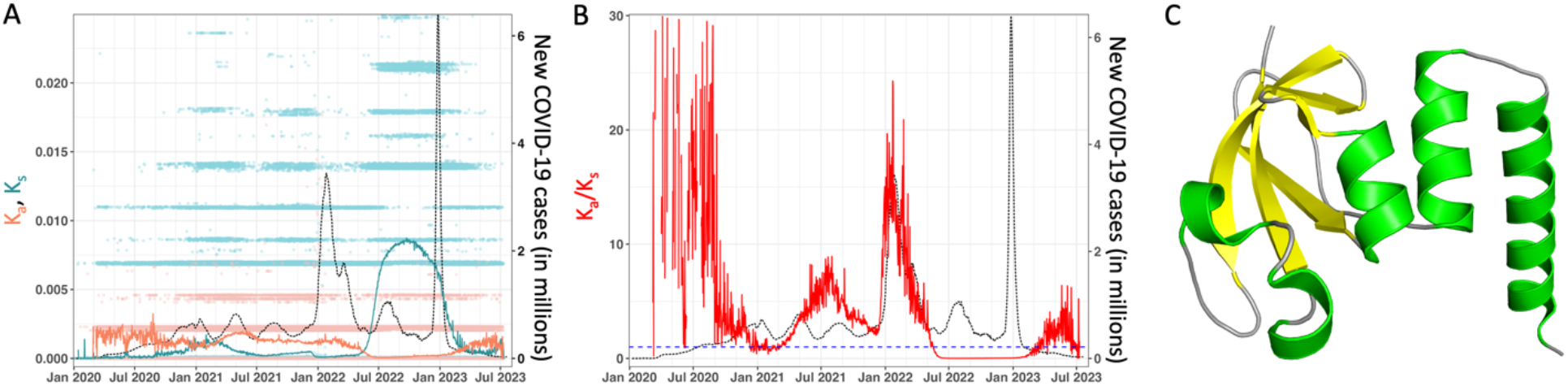
NSP8 mutation summary over the course of the COVID-19 outbreak. ((A) Non-synonymous (K_a_) and synonymous (K_s_) mutations. See Figure 5 for the VOC names corresponding to the peaks in infections and other details. (B) Ratio of non-synonymous to synonymous mutations (K_a_/K_s_). (C) Structure of NSP8, based on protein data bank (PDB) ID 7JLT.^69^

### NSP9

This protein is believed to reside in the endoplasmic reticulum.^70^ Several research efforts suggest that it interrupts host cell protein trafficking.^71^ It also is reported that NSP9 sequesters the NUP62 protein, a part of the nuclear pore complex, affecting the transportation of different components of the host’s immune response.^70^ More recently, it has been shown that this protein interacts with the E3 RING ligase enzyme, which is part of the ubiquitin system that processes different cellular functions, including immune signaling.^72^ This interaction might be part of the successful evasion of the host immune system. Like NSP7 and NSP8, amino acid substitutions have not been observed in any of the variants (Table 1). The K_s_ plot shows that the average K_s_ increases after March 2021 at the Gamma variant surge and starts to decrease again after August 2021 (Figure 17). K_s_ values dramatically increase again and stabilize at that elevated level after January 2022. These changes in K_s_ values are echoed in the nucleotide sequences in the Gamma and Omicron BA.2 – BA.5 variants.

**Figure 17:**
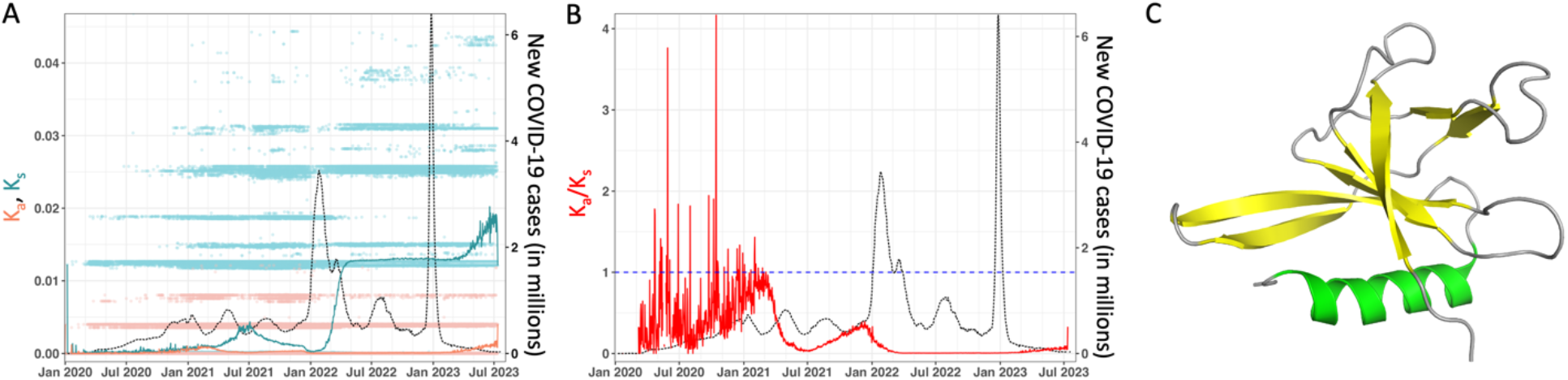
NSP9 mutation summary over the course of the COVID-19 outbreak. (A) Non-synonymous (K_a_) and synonymous (K_s_) mutations. See Figure 5 for the VOC names corresponding to the peaks in infections and other details. (B) Ratio of non-synonymous to synonymous mutations (K_a_/K_s_). (C) Structure of NSP9, based on protein data bank (PDB) ID 7BWQ.^73^

### NSP10

This protein is an essential component in activating the replication/transcription complex. It is an activator for the error-repair enzyme 3’-5’ exonuclease (NSP14) enzyme.^74,75^ Another important function attributed to this protein is its interaction with NSP16 in order to activate its 2’-O-methyltransferase activity to evade the host immune system.^75,76^ This gene has not displayed any amino acid substitutions throughout the outbreaks (Table 1). The K_s_ plot shows a significant increase around December 2021, which continues until a decrease in March 2022, coinciding with the Omicron BA.1 surge (Figure 18). This is confirmed when comparing the nucleotide alignments of the Omicron BA.1 variant against the Wuhan variant. Despite these nucleotide changes, the K_a_ plot suggests that there are no amino acid substitutions.

**Figure 18:**
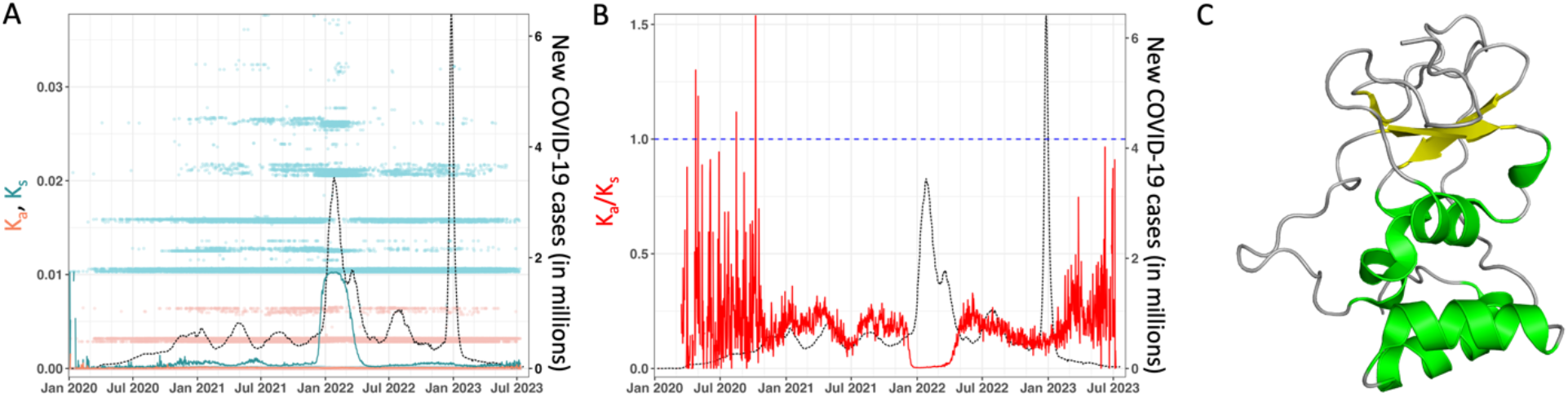
NSP10 mutation summary over the course of the COVID-19 outbreak. (A) Non-synonymous (K_a_) and synonymous (K_s_) mutations. See Figure 5 for the VOC names corresponding to the peaks in infections and other details. (B) Ratio of non-synonymous to synonymous mutations (K_a_/K_s_). (C) Structure of NSP10, based on protein data bank (PDB) ID 7MC5.^77^

### NSP11

Research suggests that this is an intrinsically disordered protein (IDP), hence it lacks an ordered structure; however, in some solvent conditions mimicking membrane environments (e.g., TFE, SDS) it forms an α-helix, suggesting that it may have some membrane-related role.^78^ As the smallest of the encoded genes, it contains just 13 amino acids. There have been no amino acid substitutions in any of the VOCs (Table 1). Due to the protein’s small size and resulting small database depth, the plots are extremely noisy; however, both individual and ratio plots show that K_a_ and K_s_ have been stable throughout all the surges (Figure 19).

**Figure 19:**
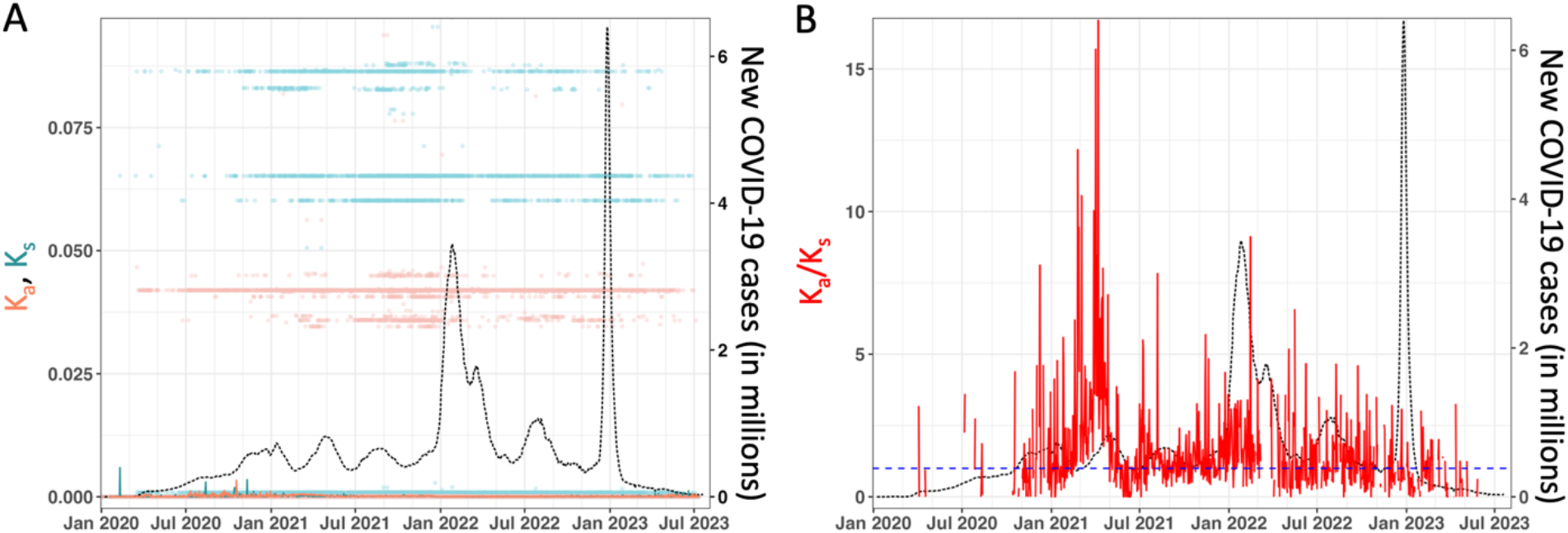
NSP11 mutation summary over the course of the COVID-19 outbreak. (A) Non-synonymous (K_a_) and synonymous (K_s_) mutations. See Figure 5 for the VOC names corresponding to the peaks in infections and other details. (B) Ratio of non-synonymous to synonymous mutations (K_a_/K_s_).

### NSP12

This is the catalytic subunit of the RNA-dependent RNA polymerase (RDRP). This catalytic subunit complexes with one molecule of NSP7 and two molecules of NSP8 and is responsible for the replication of the viral genome.^68^ Recently, a study suggested the use of modified RNA templates as a drug to inhibit NSP12.^79^

All variants share a P323L mutation when compared to the Wuhan genome (Figure 17 and Table 1). The P323L substitution is located in the RDRP interface domain. This mutation increases the stability of the RDRP structure, while simultaneously weakening the RDRP-NSP8 interaction.^80^ It is suggested that both stabilizing the RDRP structure and destabilizing its interaction with the co-factors reduces the proofreading capability, significantly increasing the RNA replication efficiency along with error-derived mutation rate.

The K_a_/K_s_ plot echoes these observations (Figure 20B). Note that the plot decreases during the Gamma variant surge between mid-April to mid-May of 2021, then immediately increases again at the Delta variant surge between June and November of 2021. It dramatically decreases again at the Omicron variant surges. While the nonsynonymous mutations are somewhat limited in scope and diversity, there are synonymous substitutions throughout the variants.

**Figure 20:**
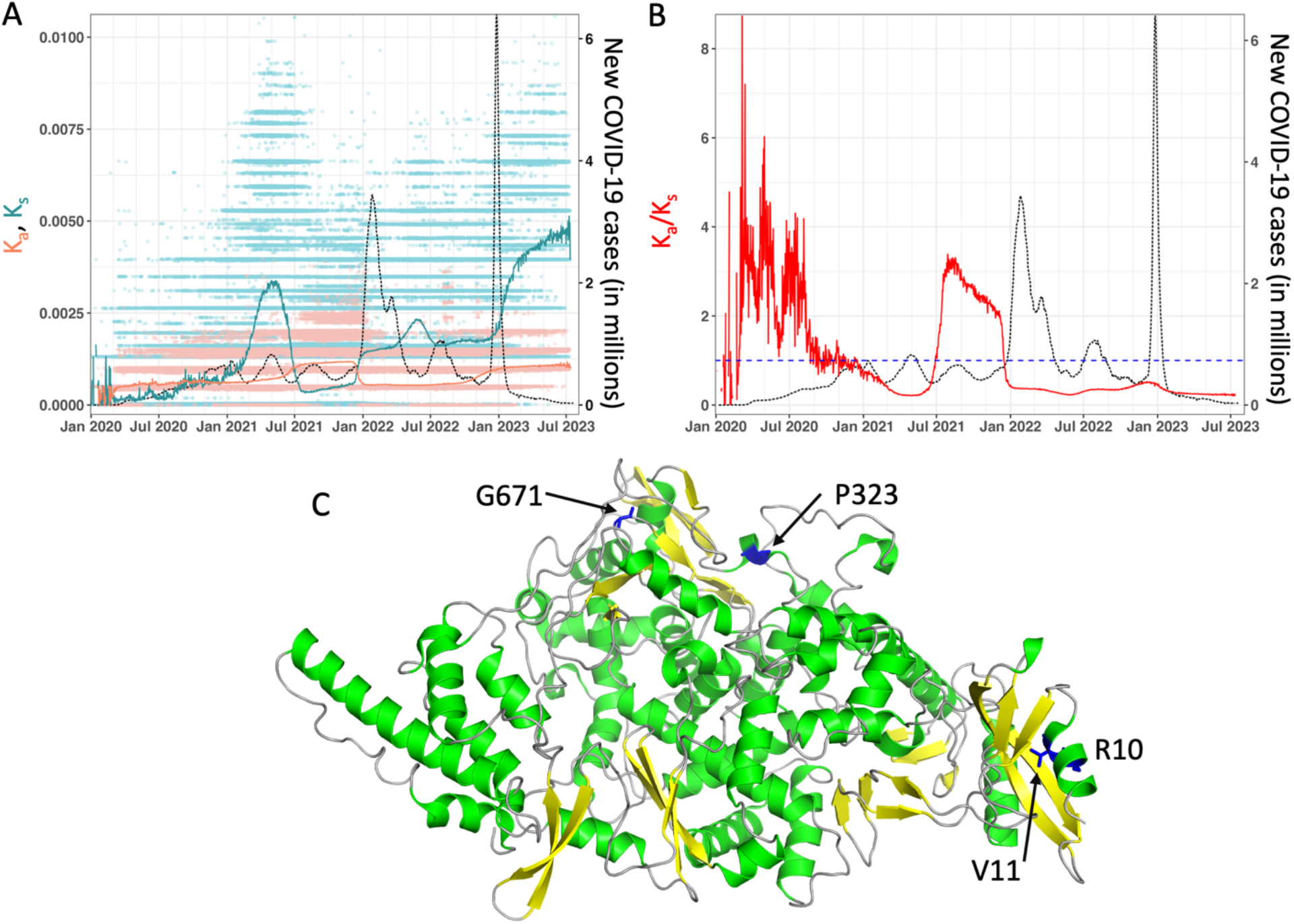
NSP12 mutation summary over the course of the COVID-19 outbreak. (A) Non-synonymous (K_a_) and synonymous (K_s_) mutations. See Figure 5 for the VOC names corresponding to the peaks in infections and other details. (B) Ratio of non-synonymous to synonymous mutations (K_a_/K_s_). (C) Structure of NSP12, based on protein data bank (PDB) ID 7KRN.^81^

### NSP13

This protein is the helicase part of the mature replication complex.^82^ There are a couple of roles attributed to this protein. One role is for proper folding of RNA genome, while the second role is as a potential suppressor of interferon I productions.^82^

Amino acid substitutions occur in all but the Alpha, Beta, and Omicron BA.1 variants, with the Omicron BA.2-BA.5 variants sharing the same R392C substitution (Figure 21 and Table 1). In accordance with molecular dynamics simulations, the Gamma variant-limited E341D mutation and Delta variant-limited P77L mutations demonstrate a higher binding affinity to the TBK1 than the Wuhan strain, thus more effectively evading the immune response.^83^ The shared Omicron variants’ mutation, R392C, is located close to the active site of the Rec1A domain. Although the effect on efficiency is yet to be determined, this mutation increases the flexibility of the protein, thus decreasing the stability.^84^ The K_a_ and K_a_/K_s_ plots reflect these nonsynonymous mutation observations (Figure 21 A-B). The average values of the K_a_/K_s_ ratio increase around the Gamma and Delta variant surges between March and September of 2021. It subsequently decreases again, until the Omicron BA.2 variant surge emerges around March of 2022. It has stayed elevated since then.

**Figure 21:**
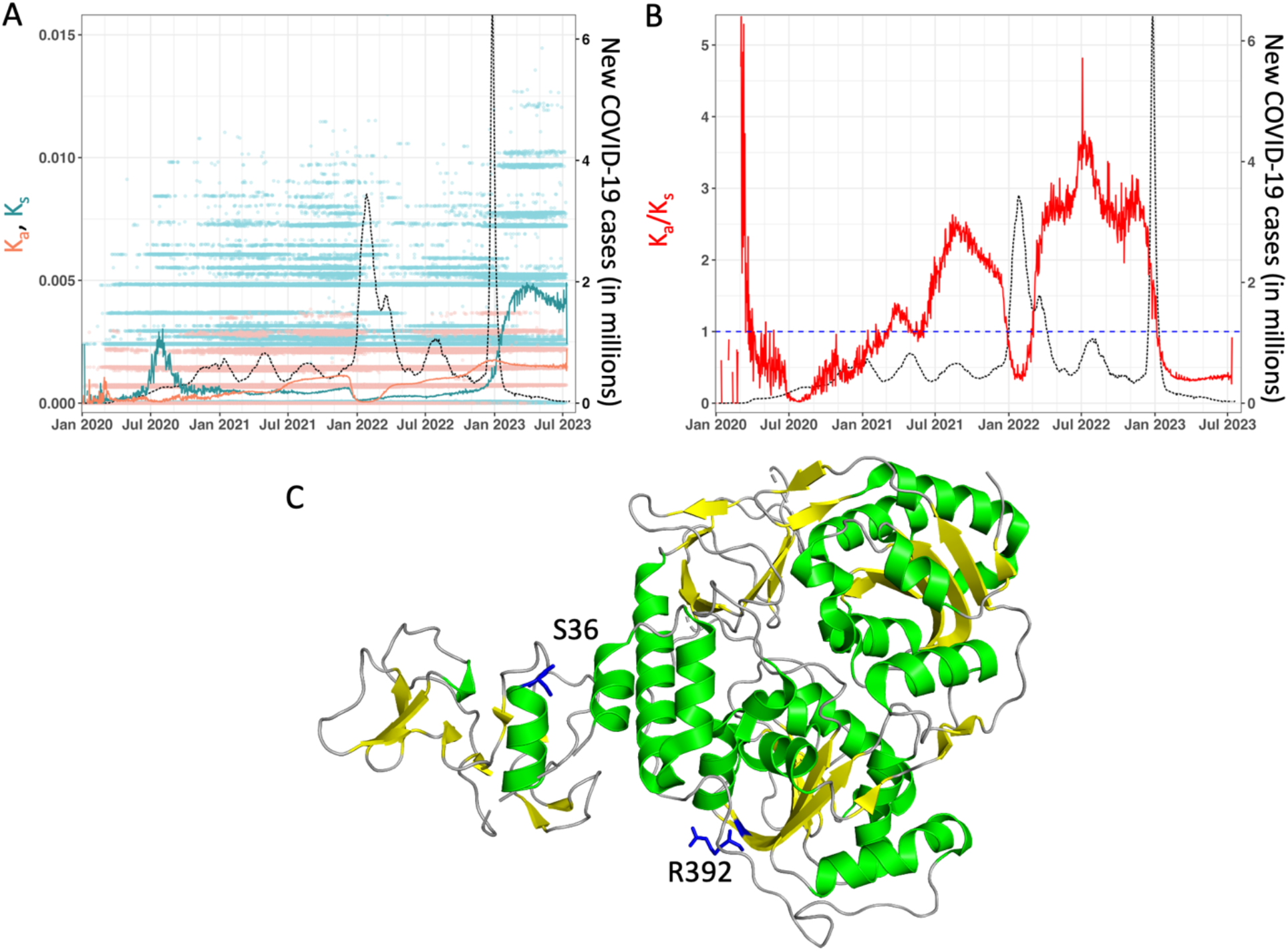
NSP13 mutation summary over the course of the COVID-19 outbreak. (A) Non-synonymous (K_a_) and synonymous (K_s_) mutations. See Figure 5 for the VOC names corresponding to the peaks in infections and other details. (B) Ratio of non-synonymous to synonymous mutations (K_a_/K_s_). (C) Structure of NSP13, based on protein data bank (PDB) ID 5RL6.^85^ Note, the missing regions in the structure are indicated by dashed lines.

### NSP14

This protein is also part of the replication complex, where it acts as the 3’ to 5’ exonuclease part of the complex.^86^ It ensures an accurate viral transcription by excising mismatched bases. Additionally, this enzyme has a second activity, which is as a C-terminal N7-methyltransferase.^87^ One of the clinically relevant roles attributed to NSP14 is interferon I response inhibition.^88^ The protein contains two unique substitutions, one of which is shared by the Omicron BA.1, BA.2, and BA.4 variants (Figure 22 and Table 1). Interestingly, all the observed substitutions are conserved hydrophobic substitutions. The K_a_/K_s_ plot shows that there is an increase in nonsynonymous amino acids substitutions starting at the Delta variant surge around June of 2021. The plot shows a continuous increase during the span of the entire Omicron subvariants surges.

**Figure 22:**
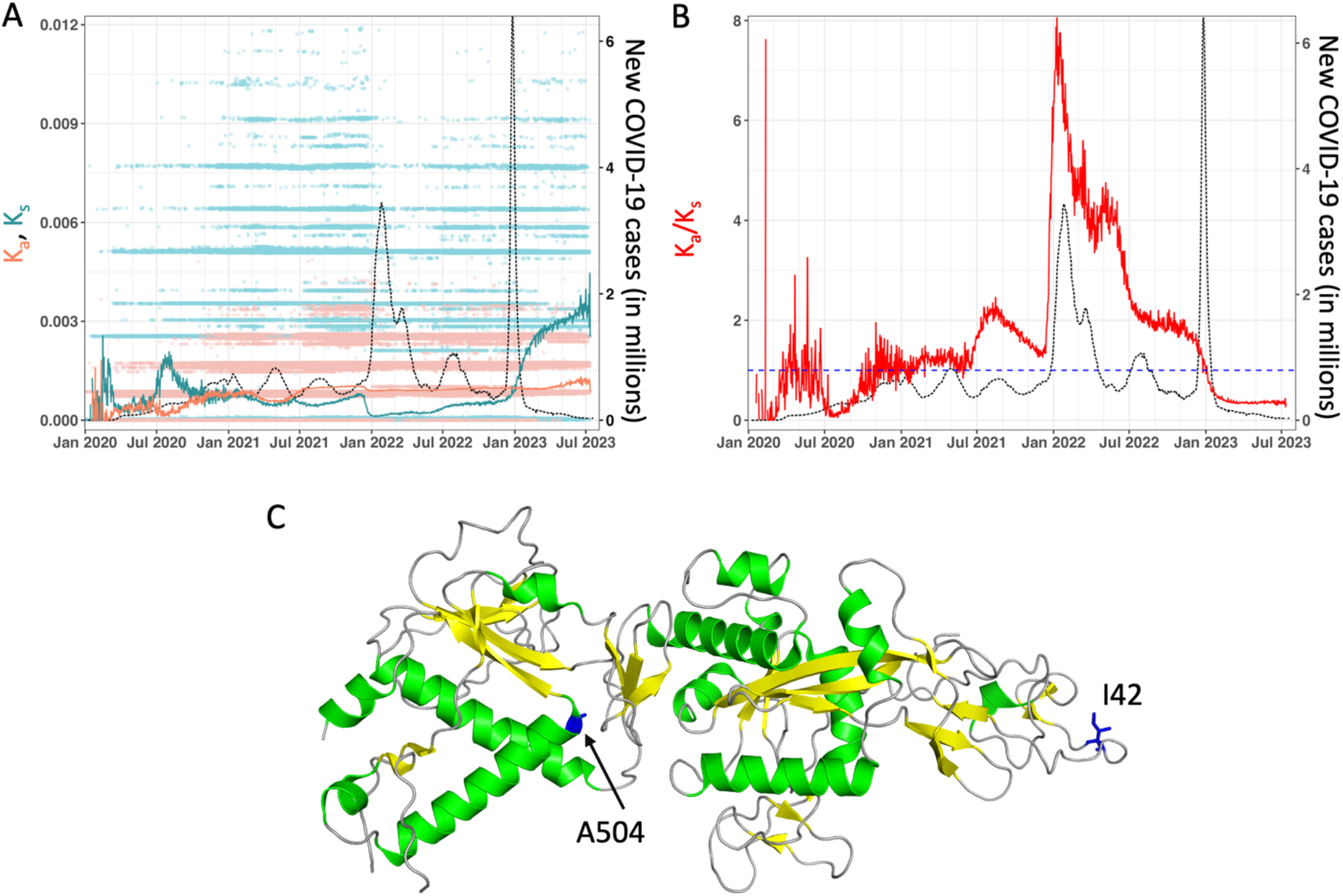
NSP14 mutation summary over the course of the COVID-19 outbreak. (A) Non-synonymous (K_a_) and synonymous (K_s_) mutations. See Figure 5 for the VOC names corresponding to the peaks in infections and other details. (B) Ratio of non-synonymous to synonymous mutations (K_a_/K_s_). (C) Structure of NSP14, based on protein data bank (PDB) ID 7N0D.^89^

### NSP15

This protein is a conserved endoribonuclease that degrades pyrimidines at the 3’ end with specificity to uridine.^90^ It preferably degrades unpaired pyrimidines.^90^ The activity of NSP15 is affected by several different factors, including RNA secondary structure. The presence of a 2’-O-ribose methyl group in the RNA inhibits the NSP15 activity.^91^ Interestingly, NSP16, which is encoded immediately downstream of it, is predicted to be a 2’-O-ribose methyl transferase. This has led to the hypothesis that NSP16 is a regulator of NSP15.^92^ The function of NSP15 is to degrade any dsRNA intermediates, thus preventing the recognition of the viral genome inside the host cell. This ultimately aids in evasion of the host immune system by delaying the type I interferon response.^93^ More interestingly, NSP15 is unique to the *nidovirus* family, with no orthologues in humans, which makes it an attractive drug target.^94^ Recently, research has shown that NSP15 binds to the E3 ligase RNF41, which suggests a disruption of the immune system.^72^

The amino acid substitution T112I is the only mutation observed (Figure 23 and Table 1). This substitution seems to be Omicron-specific, appearing in all subvariants except BA.1 and BA.5. This substitution does not match any known vital amino acid position in this protein. The plots show little mutational activity, with the K_a_/K_s_ plot dropping its small elevation at the onset of the Omicron variant in late December 2021. After the beginning of the Omicron variant surges beginning in 2022 both the K_a_ and K_s_ values remain at a steady elevated level.

**Figure 23:**
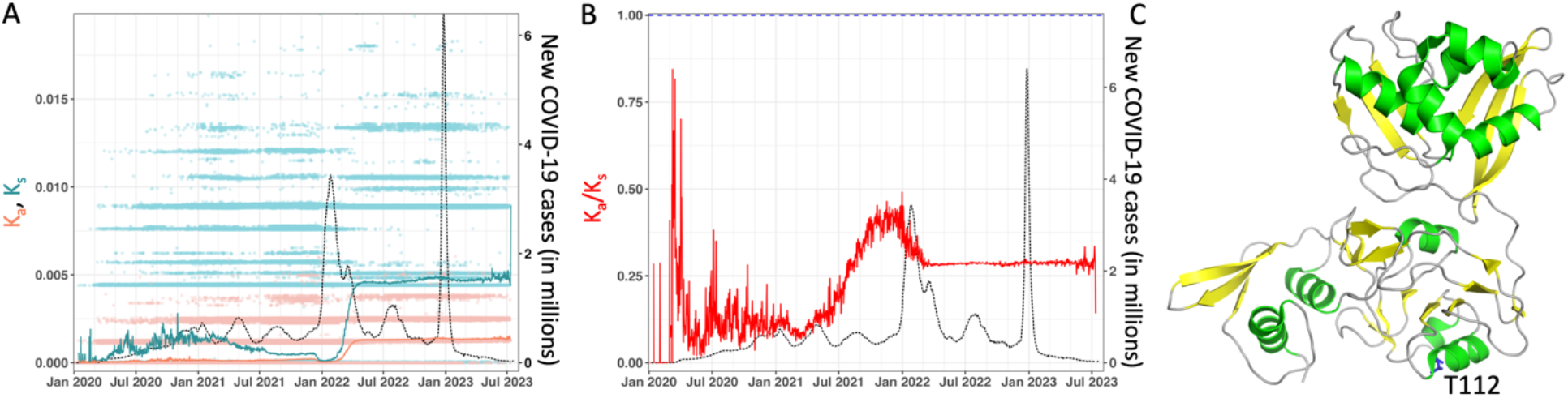
NSP15 mutation summary over the course of the COVID-19 outbreak. (A) Non-synonymous (K_a_) and synonymous (K_s_) mutations. See Figure 5 for the VOC names corresponding to the peaks in infections and other details. (B) Ratio of non-synonymous to synonymous mutations (K_a_/K_s_). (C) Structure of NSP15, based on protein data bank (PDB) ID 7ME0 [DOI:10.2210/pdb7ME0/pdb].

### NSP16

This protein is a 2’-O-methyltransferase. It has been hypothesized that it regulates NSP15.^92^ This protein also plays an essential role in immune system evasion by mimicking the activity of its human homolog, cap-specific mRNA (nucleoside-2’-O-)-methyltransferase (CMTr1), methylating mRNA to both improve translation efficiency as well as camouflaging mRNA to allow it to remain undetected from intracellular pathogen recognition receptors.^95^ Additionally, it needs to interact with NSP10 in order to be activated.^76^

No substitutions were observed for this protein in any of the major variants (Table 1). In accordance with the sequence analysis, the K_a_, K_s_, and K_a_/K_s_ plots show no noticeable change throughout the surges (Figure 24).

**Figure 24:**
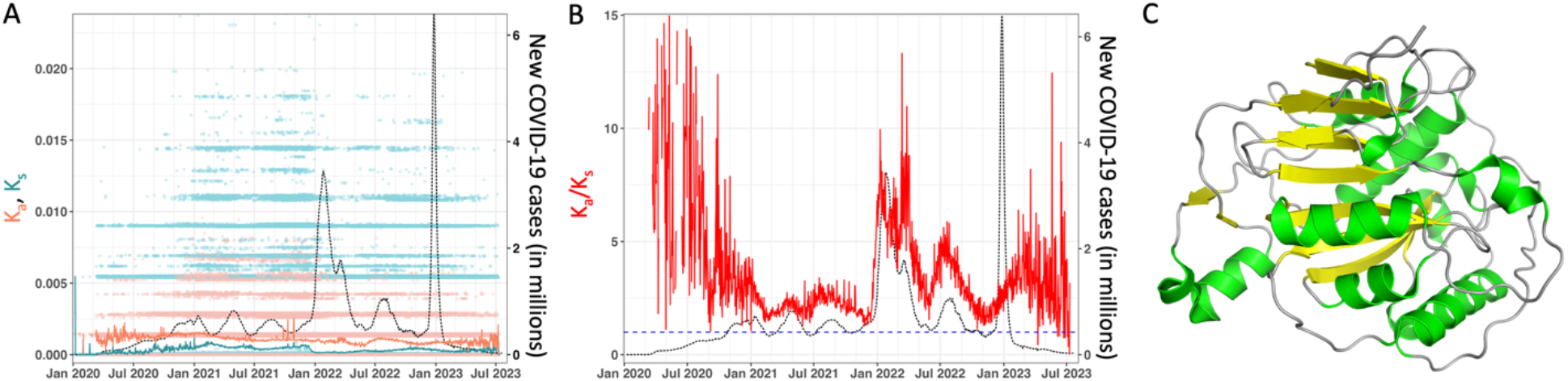
NSP16 mutation summary over the course of the COVID-19 outbreak. (A) Non-synonymous (K_a_) and synonymous (K_s_) mutations. See Figure 5 for the VOC names corresponding to the peaks in infections and other details. (B) Ratio of non-synonymous to synonymous mutations (K_a_/K_s_). (C) Structure of NSP16, based on protein data bank (PDB) ID 6WKS.^96^

### ORF3a

This protein is an integral membrane protein. It potentially is involved with the phases of the viral life cycle and viral genome replication and release.^97^ It helps to amplify viral release by promoting lysosomal exocytosis.^98^ Recently, research has shown that his protein binds to the E3 ligase TRIM59.^72^ Additionally, ORF3a, in conjunction with ORF7a, has demonstrated a reduction of antigen presentation by the human major histocompatibility complex (MHC-II), thus increasing the viral evasion of the host’s immune system.^99^

A number of different amino acid mutations occur in all variants except the Alpha and Omicron BA.1 variants (Figure 25 and Table 1). With the exception of the T223I substitution, these mutations are unique for each variant. The mutation Q57H, found in the Beta variant, was initially predicted to bind to impact binding to the spike protein, causing more severe disease;^100^ however, experimental work was found that this mutant has nearly the same activity as the Wuhan ORF3a.^101^ The Omicron BA.2 substitution L106F is located near the ion channel, although it is considered to be on the exterior side of the protein and exposed to the membrane bilayer; thus, it is unclear if the mutation has any significant impact on the protein’s function.^102^ The K_a_ plot aligns with the observed pattern and frequency of amino acids substitutions (Figure 25A). Notably, the K_a_ plot shows a sudden and steep decrease on December of 2021 corresponding to the Omicron BA.1 variant surge. Starting with the Omicron BA.2 variant surge March 2022, the plot then shows a steep increase and subsequent stability at this new level.

**Figure 25:**
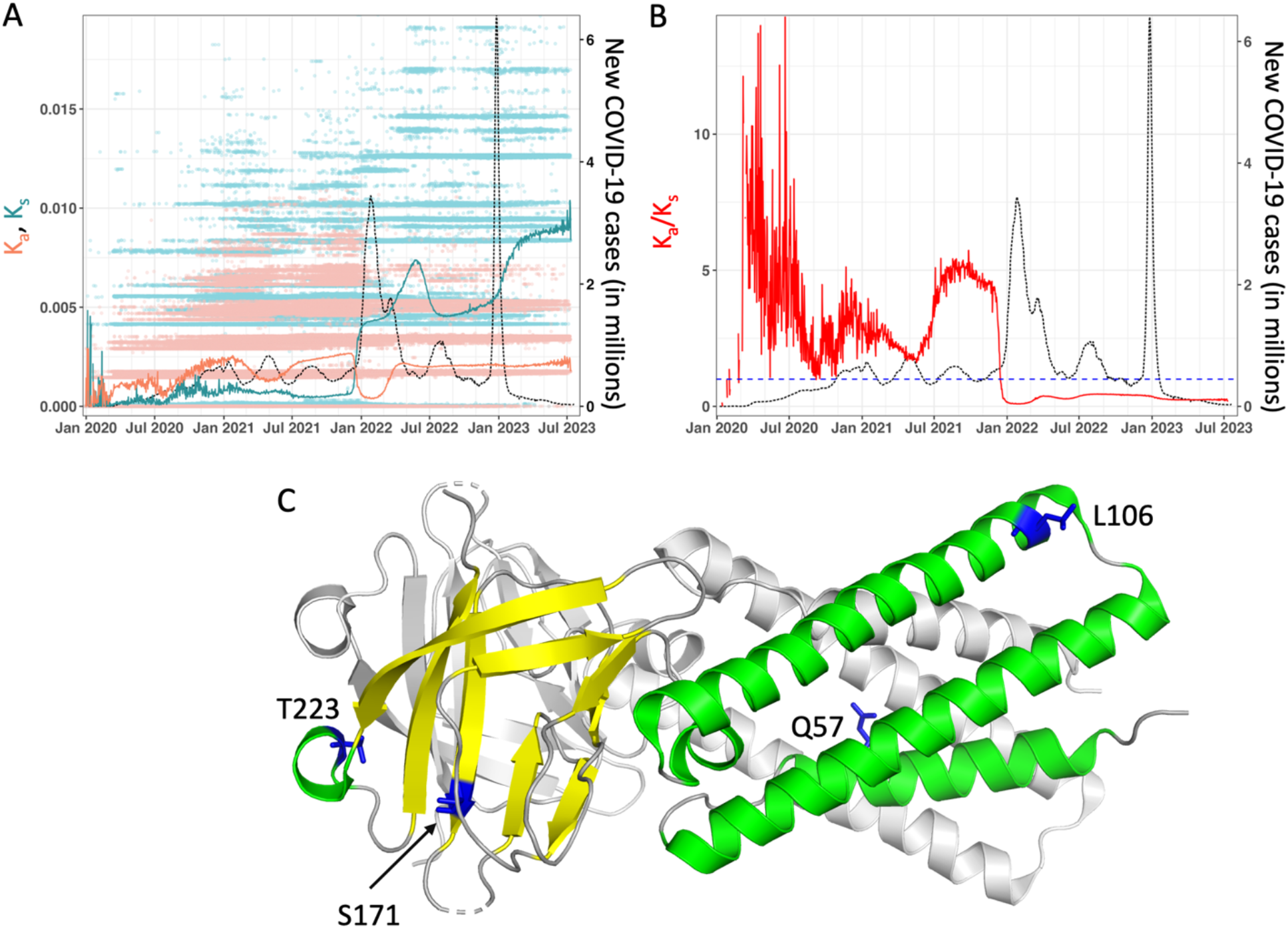
ORF3a mutation summary over the course of the COVID-19 outbreak. (A) Non-synonymous (K_a_) and synonymous (K_s_) mutations. See Figure 5 for the VOC names corresponding to the peaks in infections and other details. (B) Ratio of non-synonymous to synonymous mutations (K_a_/K_s_). (C) Structure of ORF3a, based on protein data bank (PDB) ID 6XDC.^103^ The structure is shown as a dimer, with only one protomer colored in green and yellow. The missing regions in the structure are indicated by dashed lines.

### ORF6

This protein is an antagonist of the interferon-mediated antiviral signaling pathway. Its mode of action is not well understood. However, it was demonstrated that it binds directly to the STAT1 protein resulting in its nuclear exclusion.^104^ It has also been observed that ORF6 binds the Nup98 nuclear pore component, resulting in the inhibition of the nuclear translocation of STAT1 and STAT2.^105^ More functions are speculated to be attributed to ORF6, as the localization to different membranes and its inhibitory action to the STATs are independent.^106^ We observed a D61L substitution only most Omicron subvariants, except for BA.1, BA.3, and BA.5 (Table 1). No other variants contained mutations. The amino acid substitution pattern observed is mirrored by the K_a_ plot, as the values increase in March 2022 in conjunction with the rise of the variants containing mutations (Figure 26).

**Figure 26:**
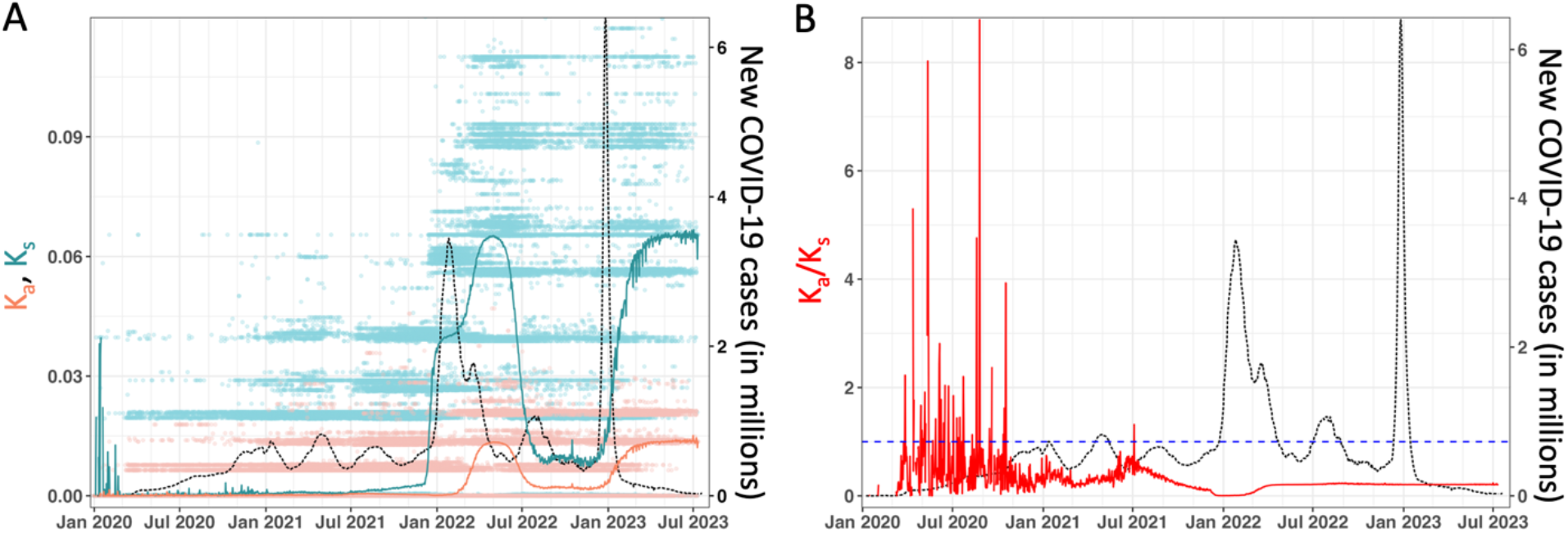
ORF6 mutation summary over the course of the COVID-19 outbreak. (A) Non-synonymous (K_a_) and synonymous (K_s_) mutations. See Figure 5 for the VOC names corresponding to the peaks in infections and other details. (B) Ratio of non-synonymous to synonymous mutations (K_a_/K_s_).

### ORF7a

This protein potentially inhibits the immobilization of the host’s tethering proteins to allow the mature viral particles to be released outside the host.^107^ It was also found to antagonize host interferon-I signaling, primarily by inhibition of STAT2 phosphorylation.^108^ In addition, a potential role in disrupting the cell cycle and inducing apoptosis has been found.^19^ Recent research has shown that overexpression of ORF7a will result in the accumulation of autophagosomes and prevention of their fusion with the lysosome, which ultimately promotes viral production.^109^ Moreover, research suggests the interaction of this protein with ORF3a to evade the immune system.^99^ The only variant that has mutations is the Delta variant with two substitutions: V82A and T120I (Figure 27 and Table 1). The change in amino acid sequence is echoed on the K_a_ plot, as the graph shows a local increase that spans the Delta variant surge and the decreases again just before the Omicron variant surges.

**Figure 27:**
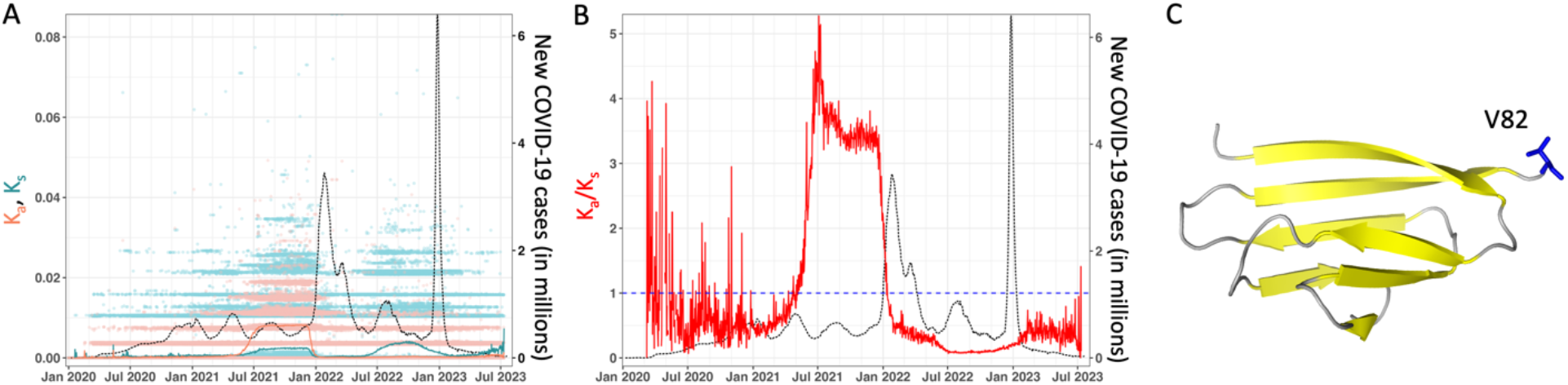
ORF7a mutation summary over the course of the COVID-19 outbreak. (A) Non-synonymous (K_a_) and synonymous (K_s_) mutations. See Figure 5 for the VOC names corresponding to the peaks in infections and other details. (B) Ratio of non-synonymous to synonymous mutations (K_a_/K_s_). (C) Structure of ORF7a, based on protein data bank (PDB) ID 7CI3.^110^

### ORF7b

This protein is believed to promote apoptosis.^111^ Additionally, it can aid in immune system avoidance through antagonism of the interferon-I signaling pathway, through the inhibition of phosphorylation of both STAT1 and STAT2.^108^ Surprisingly, the only substitution observed is L11F in the Omicron BA.4 variant (Table 1). The K_a_ plot reflects the amino acid substitution at the Delta surge (Figure 28). Immediately at the Omicron BA.1 variant surge, we can see the steep decrease of K_a_ (as well as the K_a_/K_s_ ratio) that remains stable, with only a minute increase around the Omicron BA.4 variant’s presence in June 2022.

**Figure 28:**
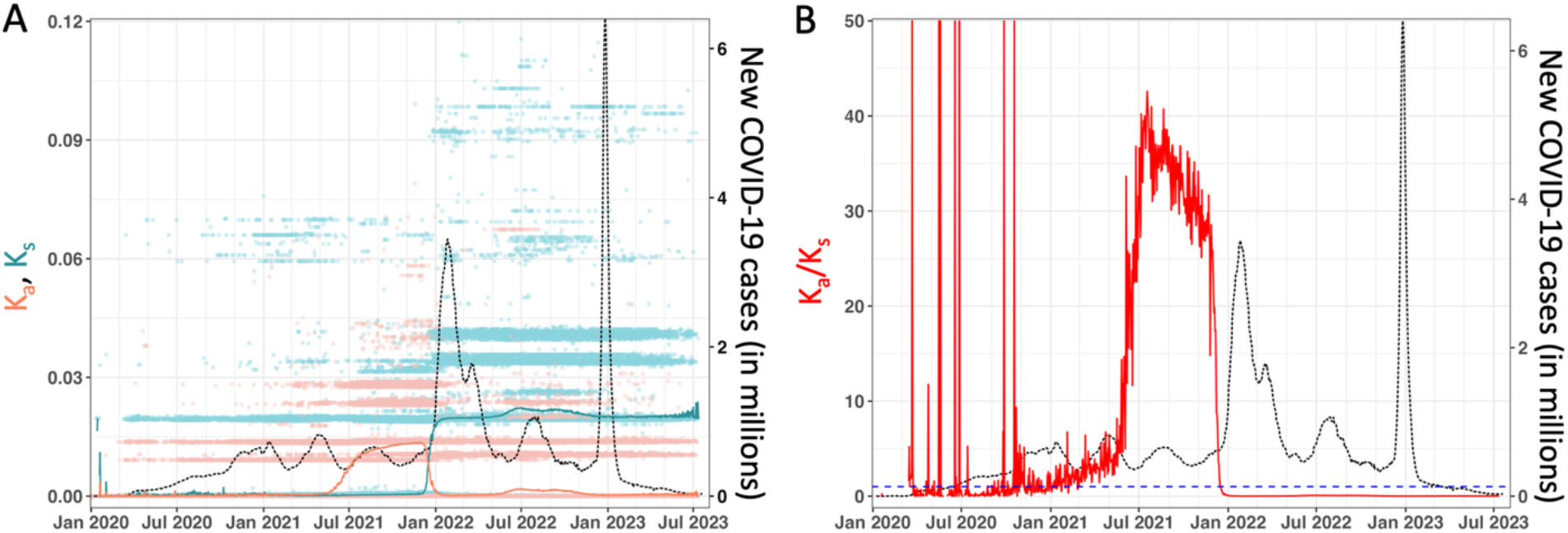
ORF7b mutation summary over the course of the COVID-19 outbreak. (A) Non-synonymous (K_a_) and synonymous (K_s_) mutations. See Figure 5 for the VOC names corresponding to the peaks in infections and other details. (B) Ratio of non-synonymous to synonymous mutations (K_a_/K_s_).

### ORF8

One of the markers for COVID-19 infection is the presence of anti-ORF8 antibodies.^112^ ORF8 has been shown to disrupt interferon-I signaling pathway and aid in immune system evasion.^112^ Sequence comparison work has found that ORF8 is a paralog of ORF7a, although it is much more dynamic and volatile than the relatively more constrained ORF7a.^113^ More recently, ORF8-knockouts have shown a noticeable decrease in lung inflammation in hamsters ^108,114^. Additionally, that same study demonstrated that a recombinant ORF8 infection would cause lymphocyte infiltration into the lung, causing severe inflammation.^114^ The two variants that have mutations pertaining to this protein are the Gamma variant with a E92K non-conservative mutation and the Delta variant with a F120L conservative mutation (Table 1). Of note is that the Alpha variant has a truncated ORF8, which seems to render it non-functional. Mutation values steadily increase until the beginning of the Omicron variant surges in December 2021, where they sharply return to 0 (Figure 29). Interestingly, synonymous mutations sharply outpace nonsynonymous ones immediately prior to the Omicron decrease.

**Figure 29:**
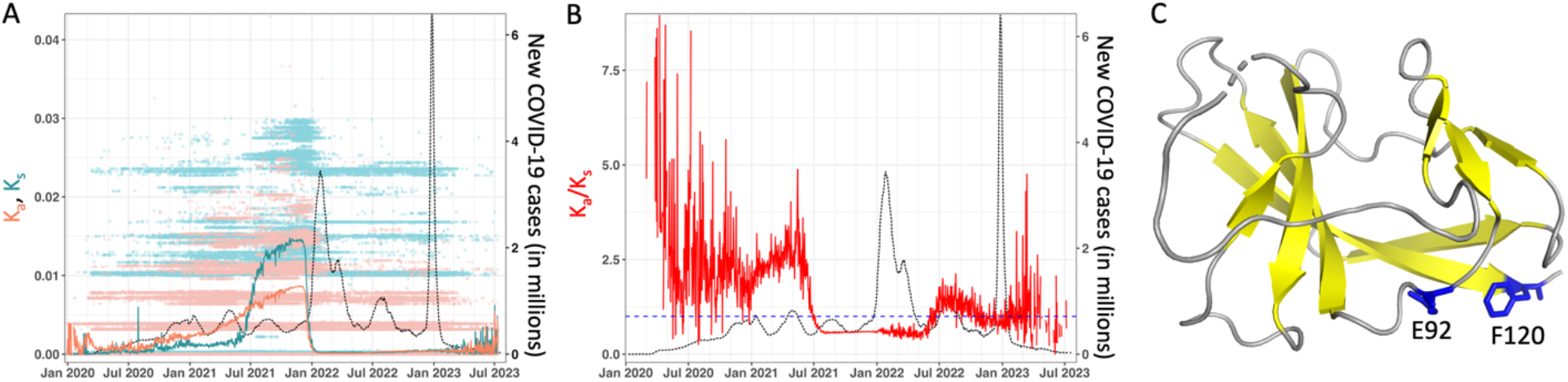
ORF8 mutation summary over the course of the COVID-19 outbreak. (A) Non-synonymous (K_a_) and synonymous (K_s_) mutations. See Figure 5 for the VOC names corresponding to the peaks in infections and other details. (B) Ratio of non-synonymous to synonymous mutations (K_a_/K_s_). (C) Structure of ORF8, based on protein data bank (PDB) ID 7JTL.^115^

### ORF10

This is a 38 amino acid-long protein. It’s been shown that it binds to E3-ubiquitin ligase, part of the host proteins degradation pathway, where the host proteins ultimately are degraded through the proteasome.^116^ This association was found to not be related to the pathogenicity of SARS-CoV-2.^116^ However, more recently, ORF10 demonstrated its ability to suppress the interferon-I signaling pathway.^117^ Specifically, the mitochondrial antiviral signaling protein (MAVS) was suggested to be the target of ORF10 suppression of the interferon-1 pathway.^117^ Our observation of this protein agrees with previous findings suggesting that this gene is under purifying pressure.^118^ We noticed only one substitution, A8V, in the Beta variant (Table 1). The paucity of VOC substitutions was reflected in the K_a_ and K_s_ graphs (Figure 30). The K_a_/K_s_ ratio graph shows extreme noise in the data, which precludes us from depicting any certain conclusions despite the apparent trend. However, it is possible that mutations were occurring outside of the dominating VOCs.

**Figure 30:**
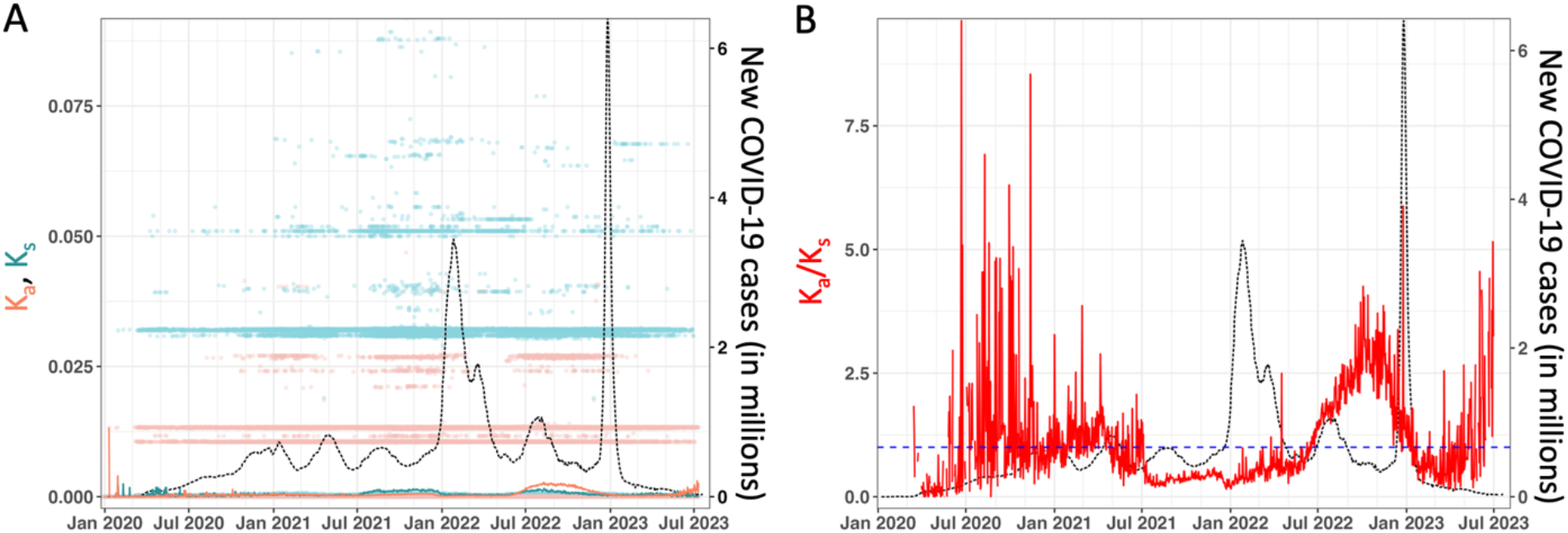
ORF10 mutation summary over the course of the COVID-19 outbreak. (A) Non-synonymous (K_a_) and synonymous (K_s_) mutations. See Figure 5 for the VOC names corresponding to the peaks in infections and other details. (B) Ratio of non-synonymous to synonymous mutations (K_a_/K_s_).

## CONCLUSIONS AND SUMMARY

Several million samples from the COVID-19 patients have now been sequenced for SARS-CoV-2 genome, and efforts are underway by the wider community to deposit these in public repositories such as NCBI’s GenBank and GISAID. As a result, an unprecedented number of SARS-CoV-2 are now available for genomic surveillance. Our group recently reported an approach to detecting mutations changes in real-time and its use for predicting the increase in number of infection cases ahead of time, with the purpose of giving medical community time for pandemic preparedness.

The analysis of over 8 million viral sequences now available in the GenBank indicate that SARS-CoV-2 continues to mutate heavily since the beginning of COVD19 outbreak. The virus is has shown a significant number of mutations, with the largest number of adaptations visible in the spike protein; most noticeably, the number of mutations have increased post-vaccination efforts. Overall, our analysis and results presented here suggest that the mutational propensities are the largest for the structural proteins, namely the spike, envelope, membrane, and the nucleocapsid proteins. Interestingly, all of these are external proteins that interact with the human immune system. On the other hand, for the majority of the period of the outbreak, the lowest number of mutations are observed in the cofactor proteins that form viral enzyme complexes. More recently, the two proteins NSP1 and NSP13 started showing an increased rate of adaptations, beginning post-Omicron BA period. New mutations have also been observed recently in NSP6, which regulates autophagy. However, NSP7-11 have not shown any significant mutations since the beginning of the pandemic.

From a genomic surveillance for pandemic surge prediction, it appears that ongoing monitoring of K_a_ provides a fairly reliable metric for surge prediction. It was noticed that for several structural proteins, the rate of mutations increased (and sometimes decreased) 2-4 weeks before the number of reported infection cases. In particular, for a number of surges in infection cases associated with various VOCs there were a number of mutations that occurred in the structural proteins as well as some NSPs. Use of the commonly used K_a_/K_s_ ratio does not appear to provide a reliable signal in most cases, even though it does tend to increase (or decrease) ahead of some surges, but there are also a significant number of cases where increase in this ratio was not followed by a surge. Based on these changes observed so far since the beginning of the pandemic, it appears that the virus continues to evolve, as it is still undergoing a significant number of adaptations. Collectively, based on the observed rates of mutations associated with relevant surges of several VOCs it is possible to make predictions about adaptations that would likely occur in future VOCs (see methods section for details). Table 2 shows a list of mutations that were predicted based on our approach on three dates, where we issued surveillance a watch or warning (full information is available on the website).

**Table 2:** Prediction of mutations in the upcoming VOC.

| Jul. 14<sup>th</sup> 2022 | Sep. 7<sup>th</sup> 2022 | Jan. 7<sup>th</sup> 2023 |
| --- | --- | --- |
| **Spike:** T19I, Δ24, Δ25, Δ26, A27S, Δ69, Δ70, G142D, V213G, G339D, S371F, S373P, S375F, T376A, D405N, R408S, K417N, N440K, L452R, S477N, T478K, E484A, F486V, Q498R, N501Y, Y505H, D614G, H655Y, N679K, P681H, N764K, D796Y, Q954H, N969K<br>**Envelope:** T9I<br>**Membrane:** D3N, Q19E, A63T<br>**Nucleocapsid:** P13L, Δ31, Δ32, Δ33, R203K, G204R, S413R<br>**NSP1:** S135R<br>**NSP4:** L264F, T327I, T492I<br>**NSP5:** P132H<br>**NSP6:** Δ105, Δ106, Δ107, F108L<br>**NSP12:** P323L<br>**NSP13:** R392C<br>**NSP14:** I42V<br>**NSP15:** T112I<br>**NSP16:** T140I<br>**ORF10:** L37F | **Spike:** T19I, Δ24, Δ25, Δ26, A27S, Δ69, Δ70, G142D, V213G, G339D, S371F, S373P, S375F, T376A, D405N, R408S, K417N, N440K, L452R, S477N, T478K, E484A, F486V, Q498R, N501Y, Y505H, D614G, H655Y, N679K, P681H, N764K, D796Y, Q954H, N969K<br>**Envelope:** T9I<br>**Membrane:** D3N, Q19E, A63T<br>**NSP1:** S135R<br>**NSP4:** L264F, T327I, T492I<br>**NSP5:** P132H<br>**NSP6:** Δ105, Δ106, Δ107, F108L<br>**NSP12:** P323L<br>**NSP13:** R392C<br>**NSP14:** I42V<br>**NSP15:** T112I<br>**NSP16:** T140I<br>**ORF3a:** T223I<br>**ORF6:** D61L<br>**ORF10:** L37F | **Spike:** L5F, M153T, N164K, H245N, G257D, K444R, N450D, L452M, N460K, E484R, G485D<br>**NSP2:** T547I<br>**NSP3:** I896T, G1273D, M1351V, T1356I, T1830I<br>**NSP4:** A146V, L438F<br>**NSP7:** Q63R<br>**NSP12:** D155G<br>**NSP13:** T481M<br>**ORF3a:** L140F |

Mutations predicted based on method described in Figure 2. See https://pandemics.okstate.edu/covid19/ for more details.

The presented framework and the real-time project website enable the tracking of the new mutations in real-time and possibly enable prediction of future variants which are relevant for diagnostic kits, as large changes could potentially make detection by the antibodies difficult. Similarly, for the vaccine and drug design efforts it would be important to keep an eye on the changes. The research community gained a wealth of knowledge about SARS-CoV-2 and its genome due to a considerable effort world-wide to tackle the pandemic, with sustained efforts to sequence millions of sequences enabling genomic surveillance efforts. Arguably, perhaps the community learned more about the genome of this virus in a shorter period of time than any other virus because of efforts across the globe to both sequence and deposit results in public databases. The recent downturn in the number of samples from COVID-19 positive patients being sequenced and being publicly being reported could undermine the genomic surveillance efforts. Therefore, the presented results serve as a call to the medical and health community to keep dedicating resources for regularly sequencing the virus. This prototype would also serve as a model for future pandemics, where the presented framework could easily be reworked and immediately be adapted from day one.

