## Supporting Information for "Pandemic preparedness through genomic surveillance: Overview of mutations in SARS-CoV-2 over the course of COVID-19 outbreak"

**Table S1: Sequence counts for SARS-CoV-2 proteins.***

| Protein | Total Sequences | Unique Nucleotide Sequences | Unique Amino Acid Sequences |
|---|---|---|---|
| NSP1 | 3235252 | 15214 | 7561 |
| NSP2 | 3190042 | 79704 | 29919 |
| NSP3 | 2649696 | 288242 | 128048 |
| NSP4 | 3129709 | 35248 | 13585 |
| NSP5 | 3412055 | 13701 | 3780 |
| NSP6 | 3290353 | 18688 | 8010 |
| NSP7 | 3556903 | 1469 | 486 |
| NSP8 | 3453121 | 5195 | 1692 |
| NSP9 | 3538898 | 3246 | 999 |
| NSP10 | 3438748 | 2729 | 783 |
| NSP11 | 3501922 | 93 | 62 |
| NSP12 (RDRP) | 3192294 | 72319 | 17102 |
| NSP13 | 3295092 | 42930 | 15314 |
| NSP14 | 2970513 | 34500 | 11129 |
| NSP15 | 3276709 | 14655 | 5954 |
| NSP16 | 3161206 | 8364 | 3287 |
| Spike | 2357986 | 227602 | 115658 |
| ORF3a | 3343136 | 47207 | 25977 |
| Envelope | 3470634 | 1626 | 894 |
| Membrane | 3120031 | 12767 | 3585 |
| ORF6 | 3512622 | 2324 | 1055 |
| ORF7a | 3242202 | 10506 | 5301 |
| ORF7b | 3298939 | 1489 | 793 |
| ORF8 | 2222565 | 7700 | 4959 |
| Nucleocapsid | 3258161 | 81663 | 34372 |
| ORF10 | 3338877 | 756 | 418 |

Last Updated: 2023-07-27 09:04:00 (Central Time)
Latest Collection Date: 2023-07-20
Total records:
Accession Count: 3596381 (quality controlled)

*Only protein sequences noted as complete and lacking ambiguous bases have been included in our dataset. Accessions without one or more qualifying protein sequence are not included in these quality-controlled sequences (listed below).

**Alternative $K_a/K_s$' calculation**

As described in the main manuscript, an alternate method to calculate the $K_a/K_s$ values was used. (see method 1 described in the Methods section for details). The results plots for the 26 proteins are shown below. Note, the individual $K_a/K_s$' values are shown as pink dots, with the red curve showing the weighted daily mean values. The dotted curve shows new COVID-19 cases (sliding weekly average) across the world. A blue dashed line denotes a value of 1; for the explanation of the significance of this value, see the main text.

**Figure S1: NSP1**

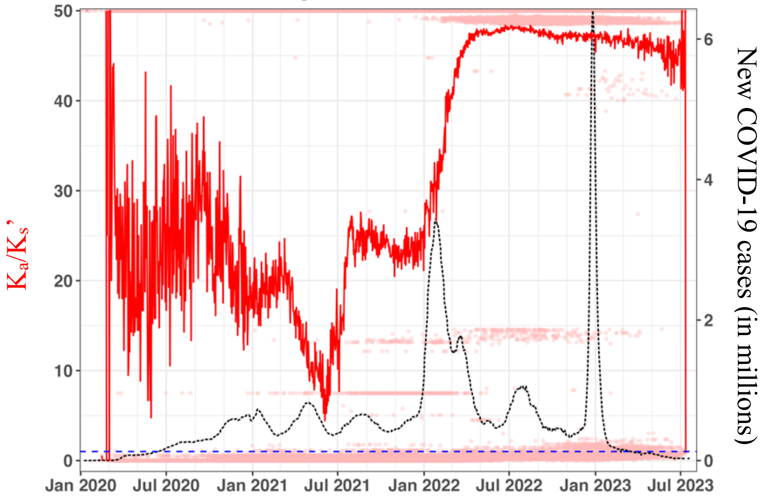


**Figure S2: NSP2**

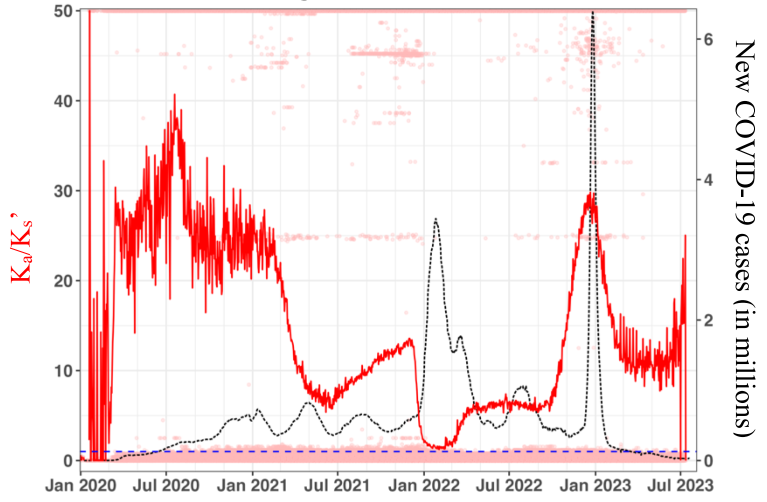

### Figure S3: NSP3

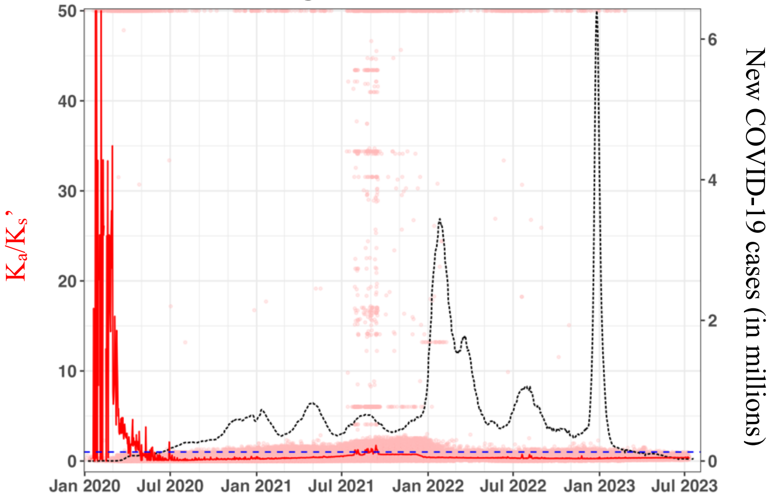


### Figure S4: NSP4

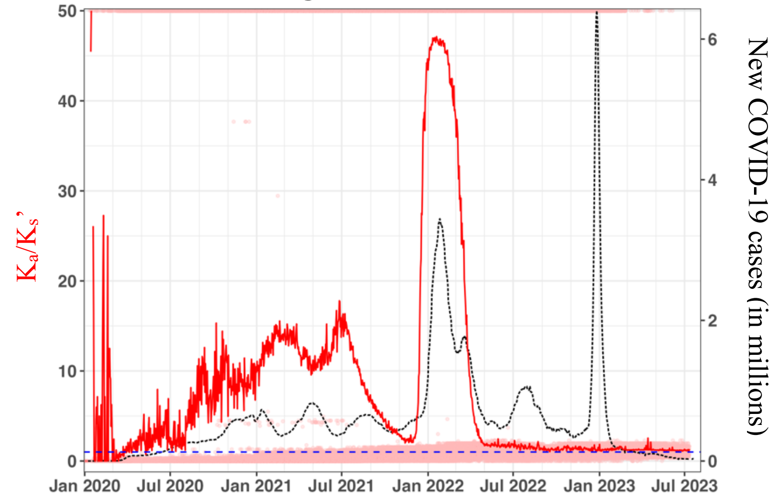


### Figure S5: NSP5

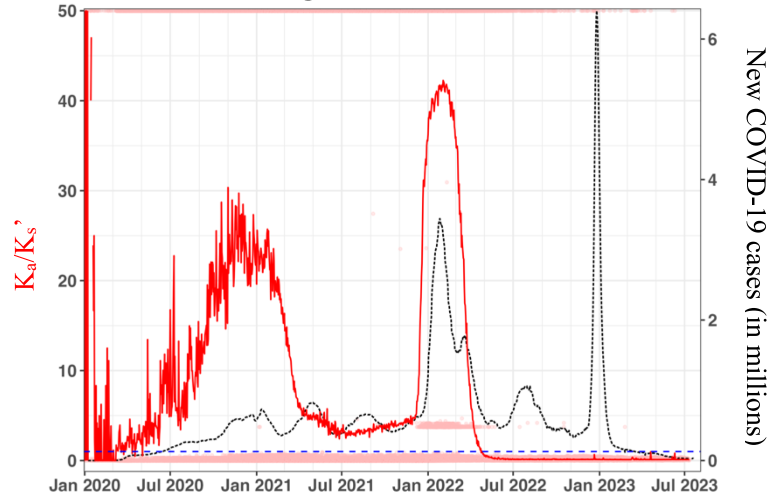

**Figure S6: NSP6**

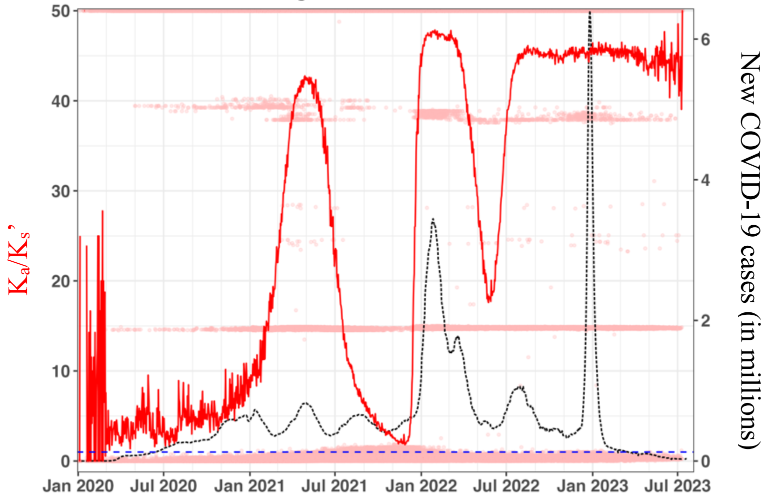


**Figure S7: NSP7**

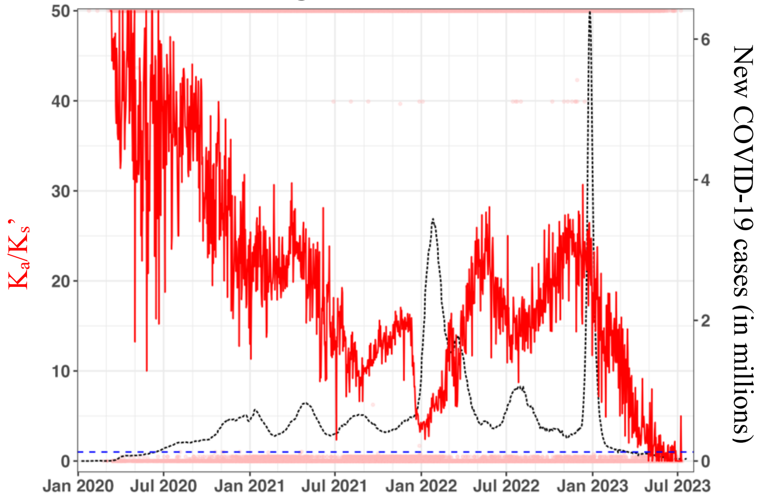


**Figure S8: NSP8**

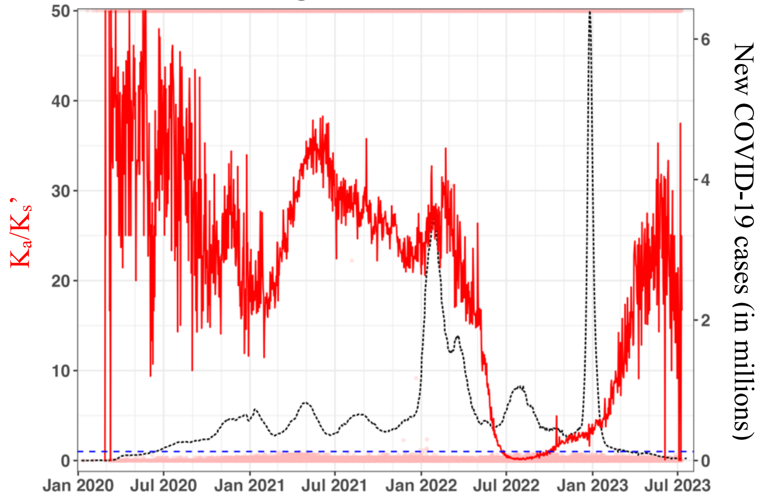

### Figure S9: NSP9

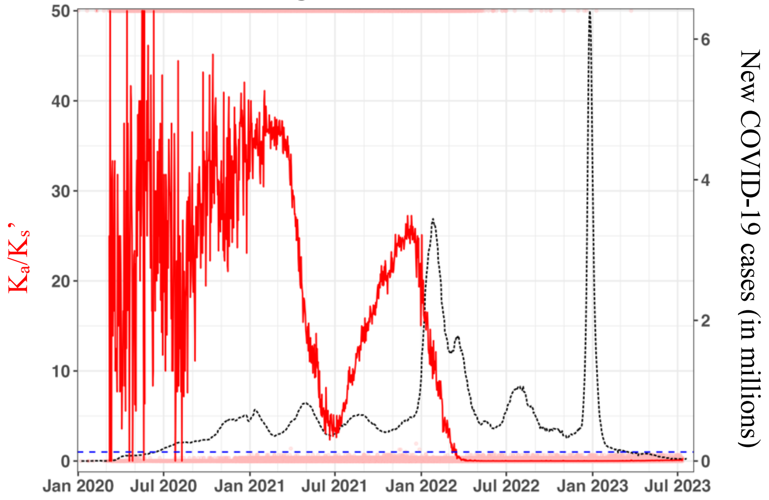


### Figure S10: NSP10

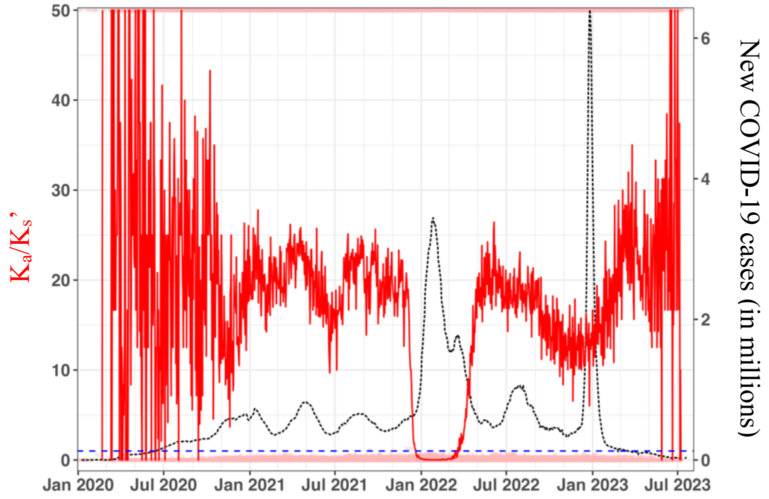


### Figure S11: NSP11

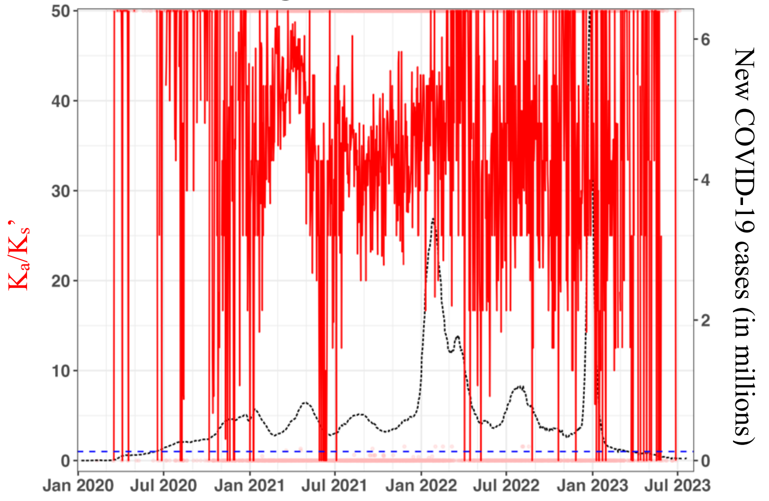

### Figure S12: NSP12 (RDRP)

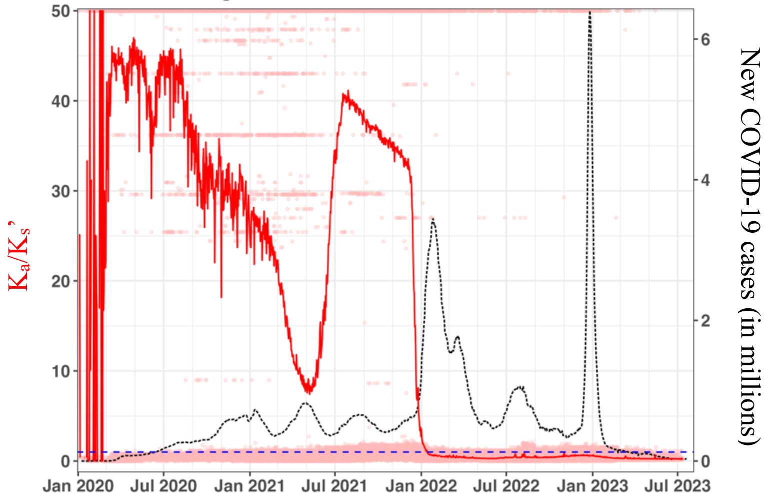


### Figure S13: NSP13

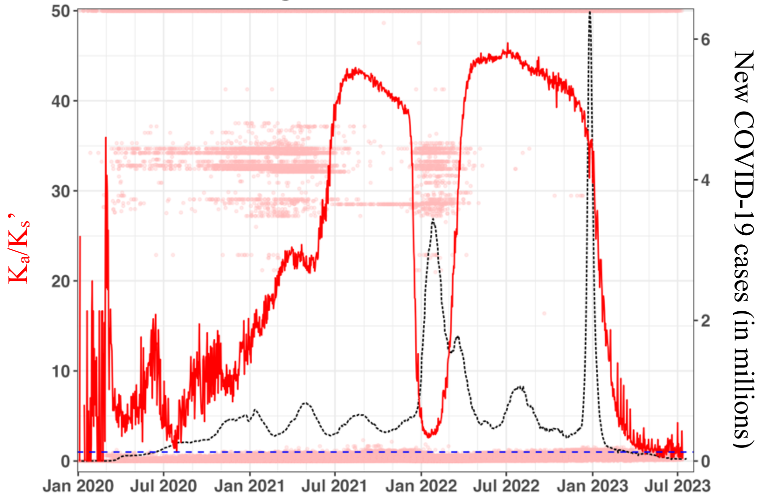


### Figure S14: NSP14

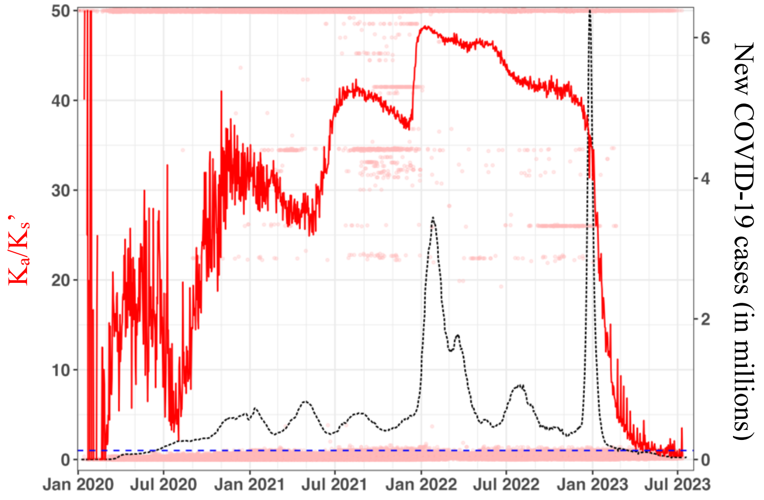

### Figure S15: NSP15

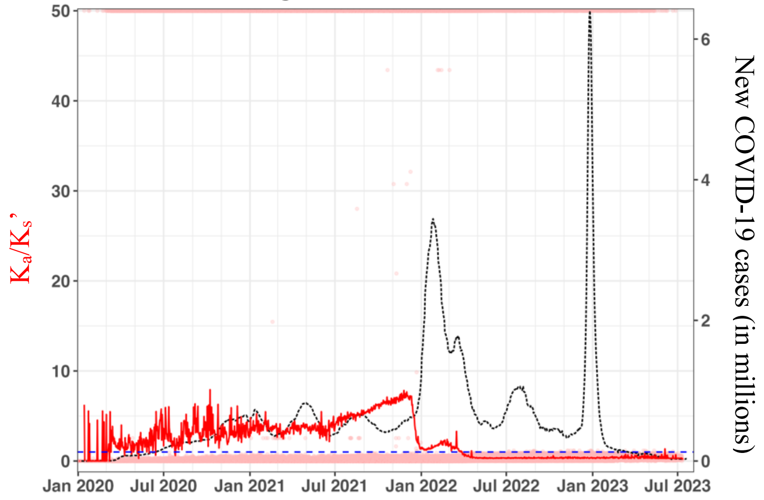


### Figure S16: NSP16

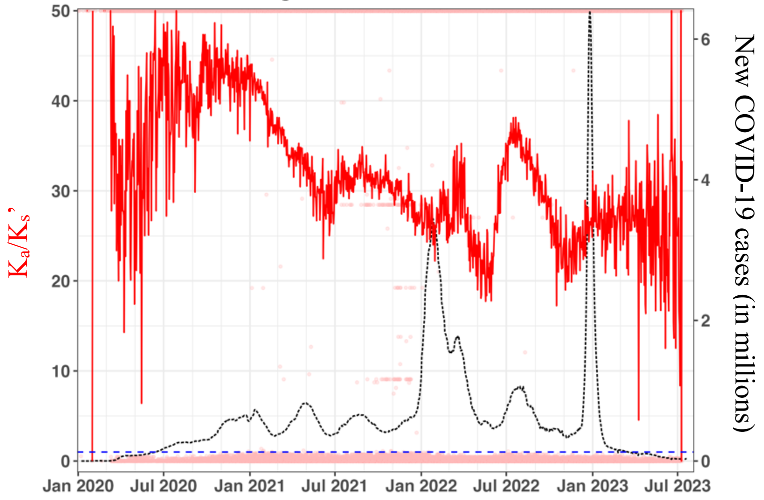


### Figure S17: Spike protein

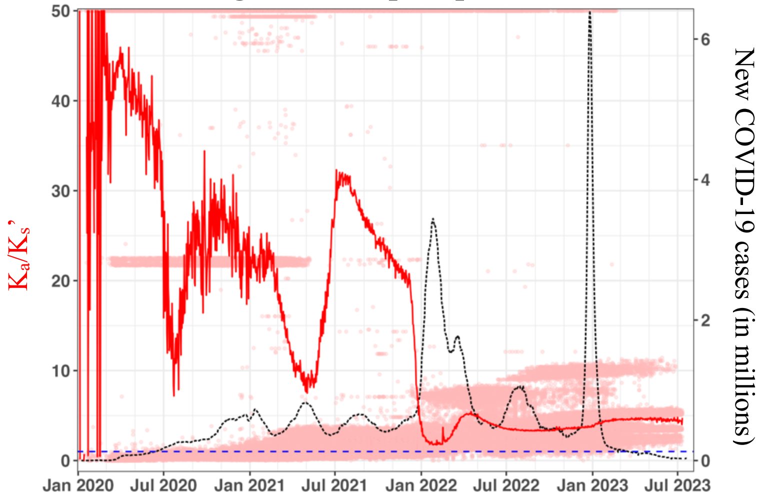

### Figure S18: Envelope protein

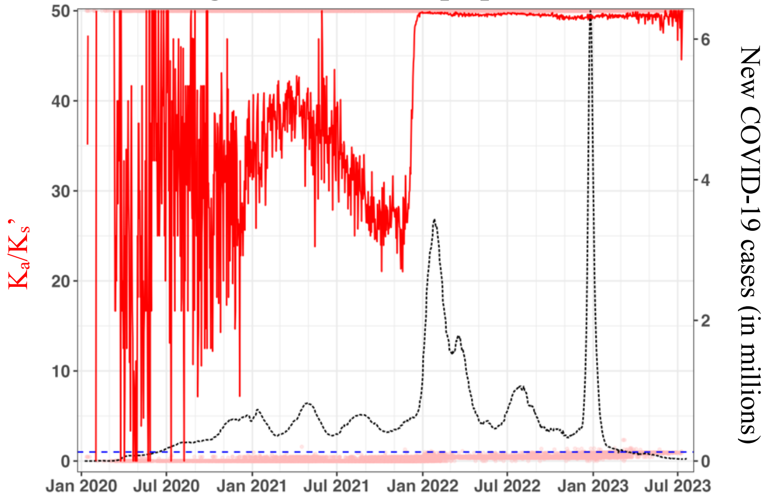


### Figure S19: Membrane protein

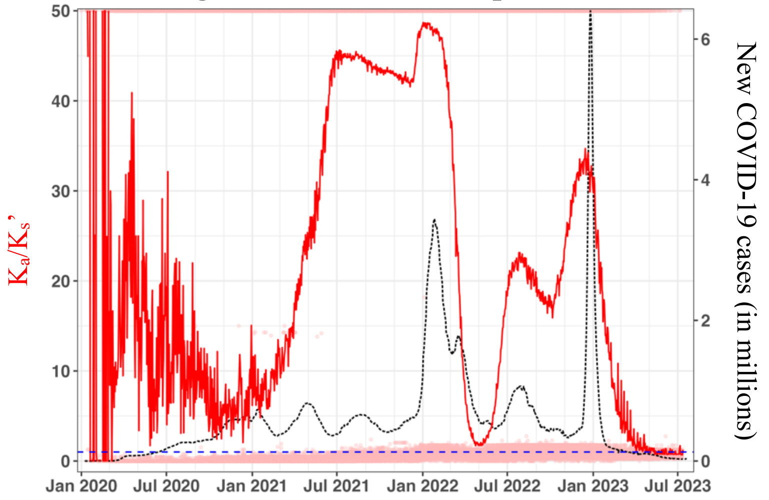


### Figure S20: Nucleocapsid protein

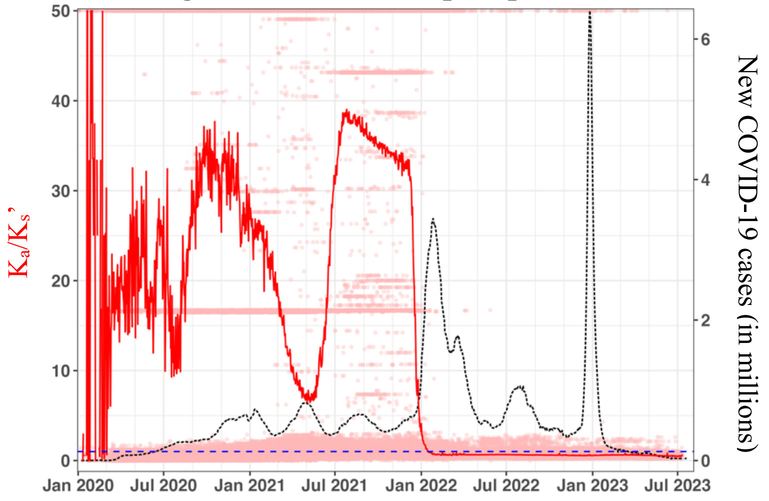

### Figure S21: ORF3a

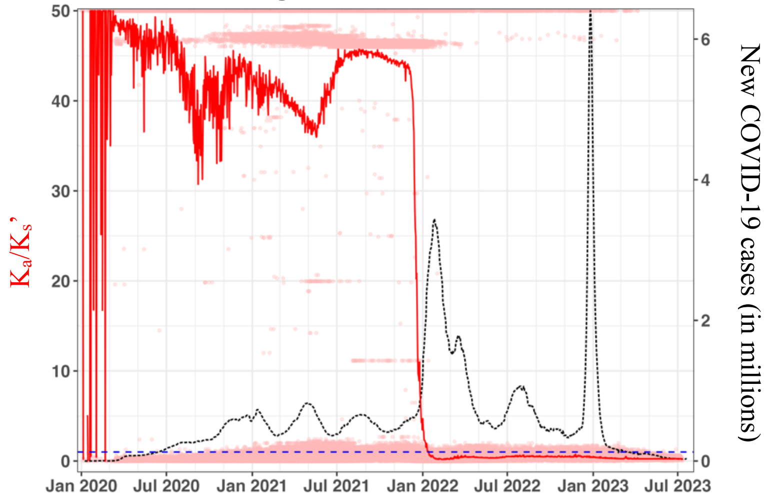


### Figure S22: ORF6

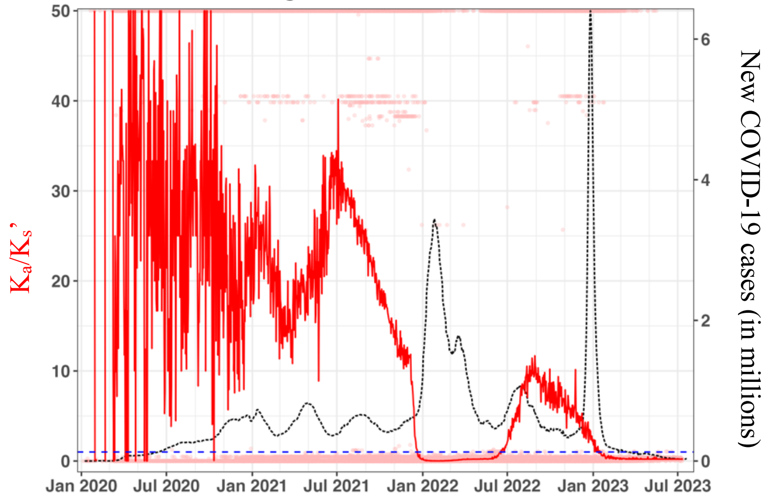


### Figure S23: ORF7a

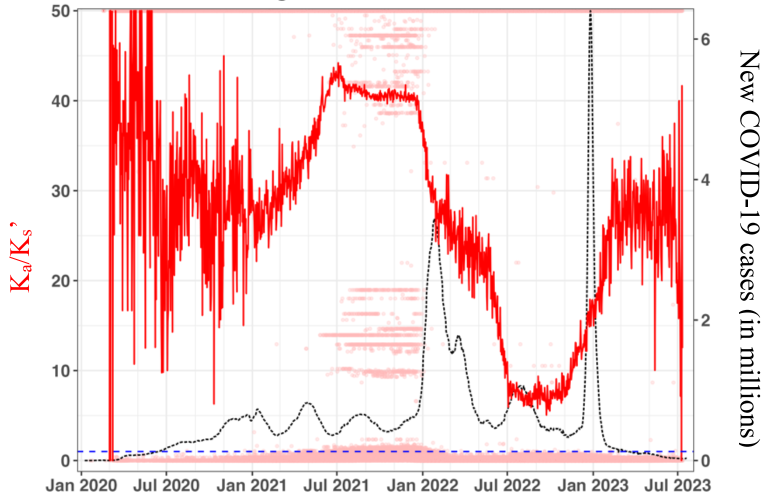

### Figure S24: ORF7b

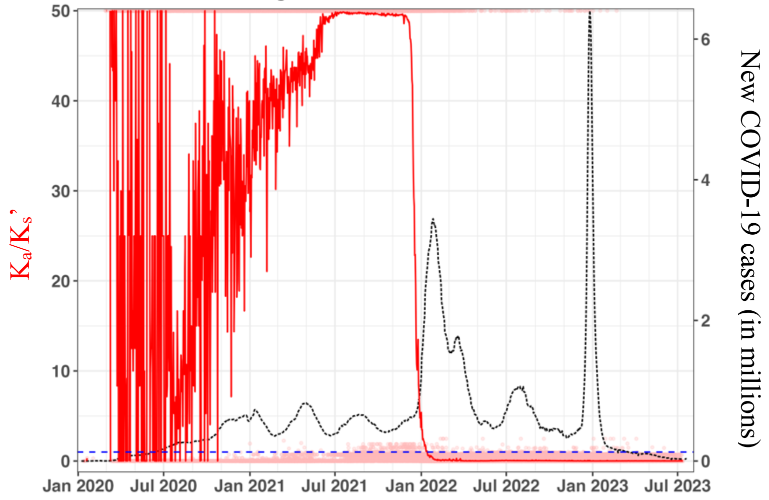


### Figure S25: ORF8

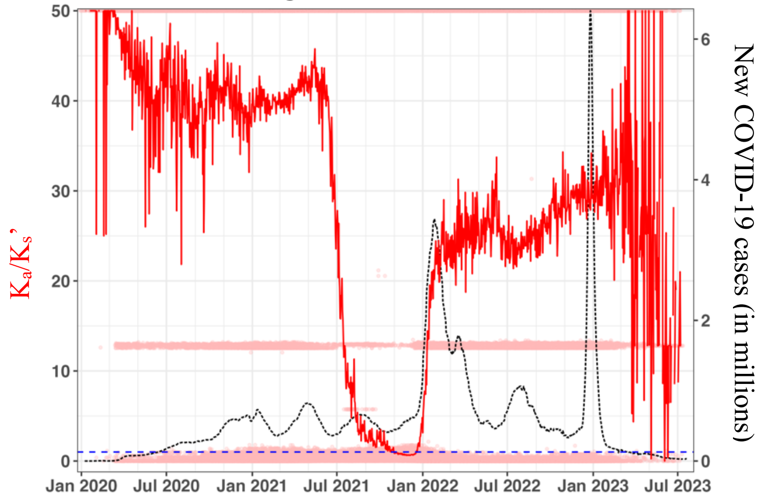


### Figure 26: ORF10

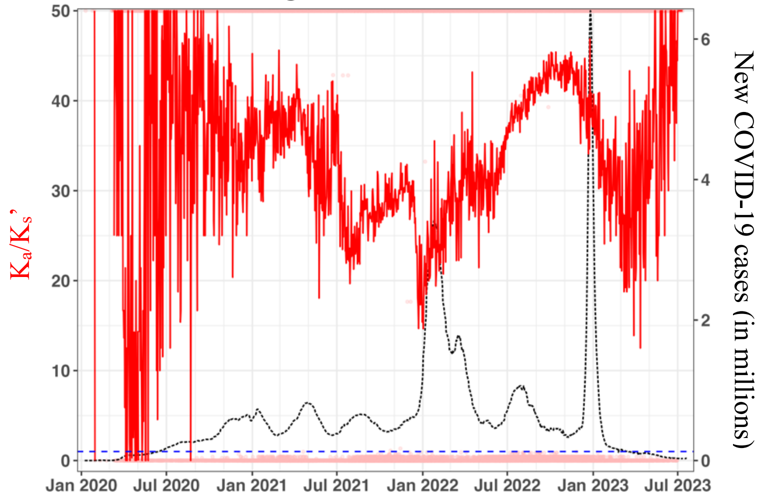

**Figure S27: $K_a/K_s$ of VOCs**

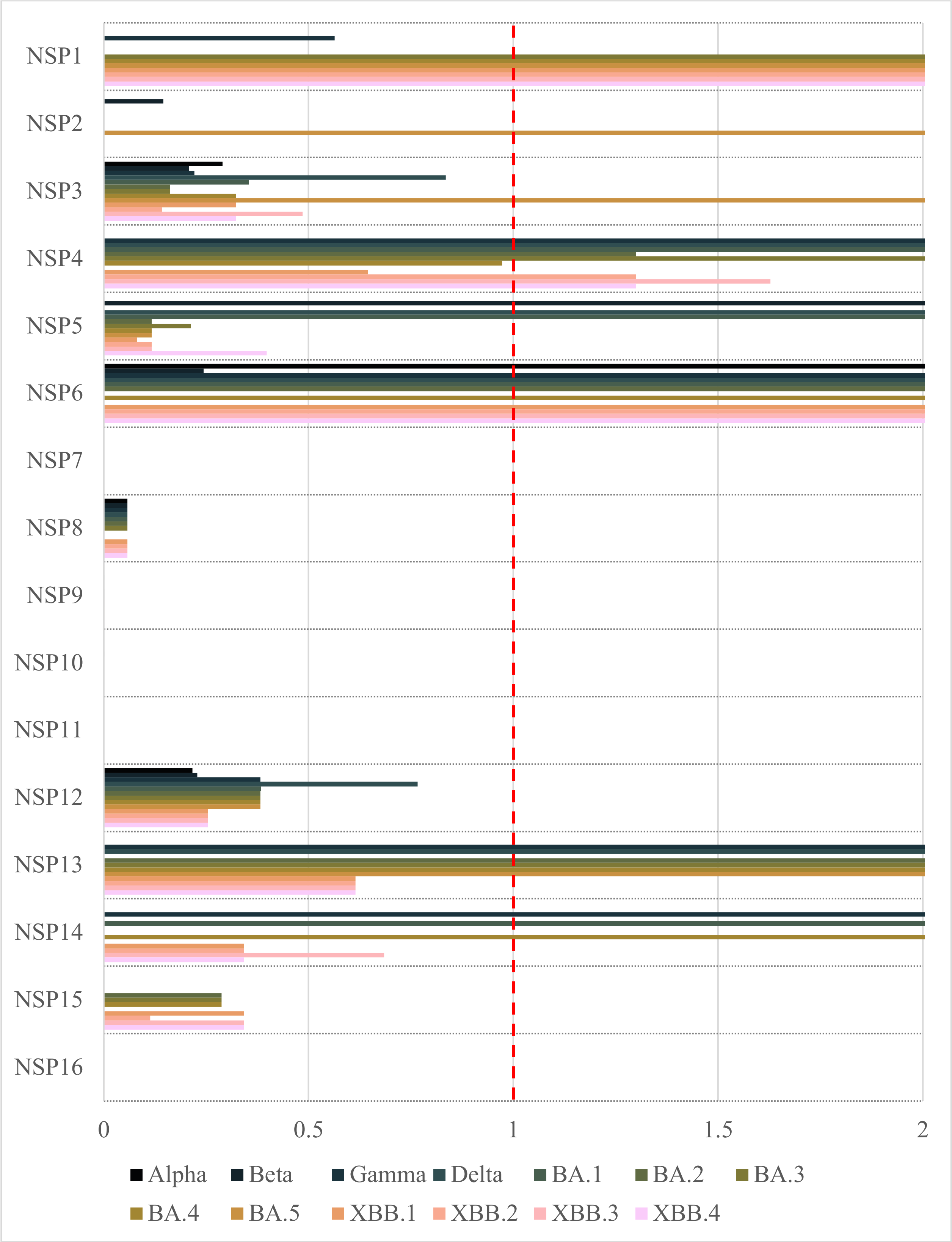

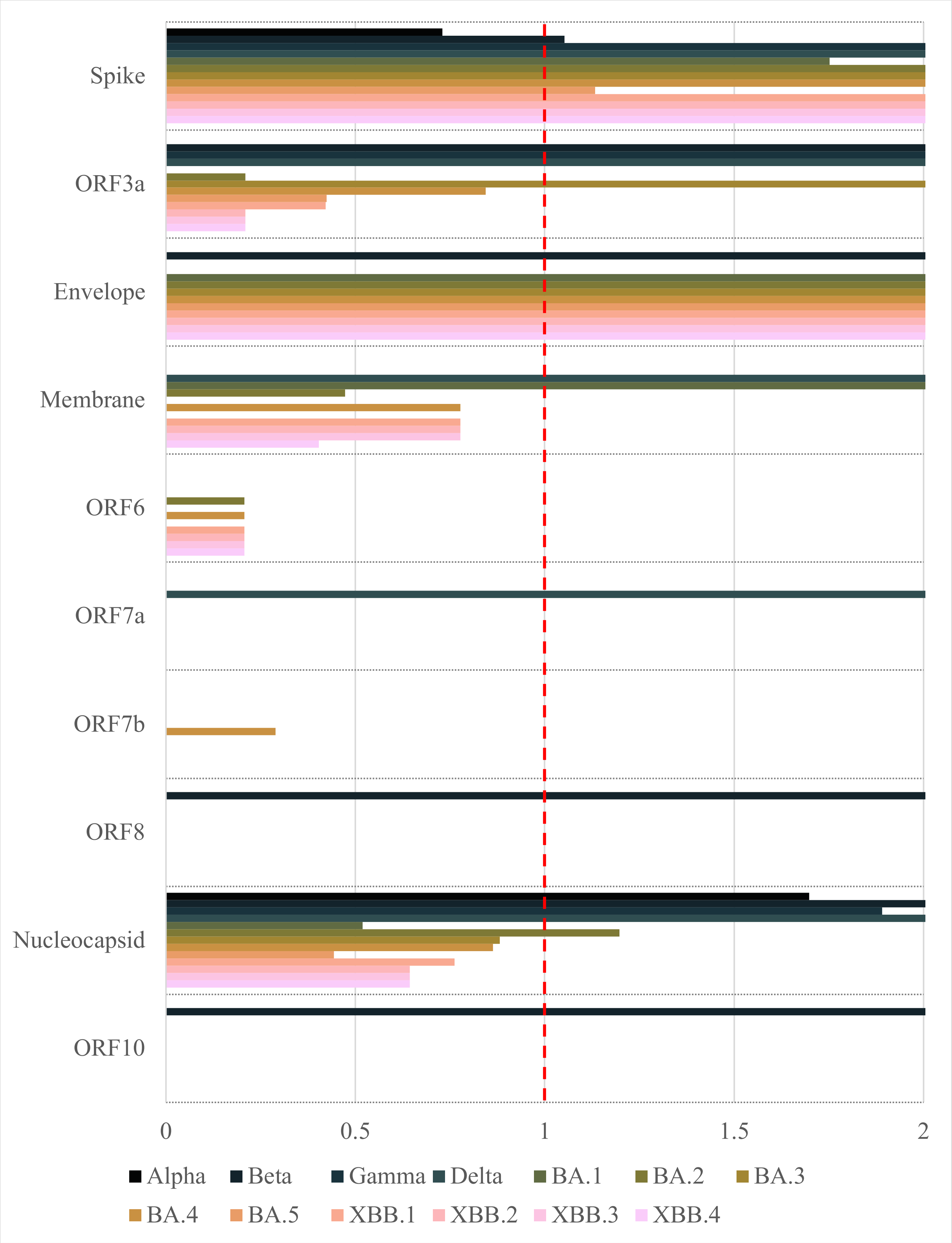

Spike
ORF3a
Envelope
Membrane
ORF6
ORF7a
ORF7b
ORF8
Nucleocapsid
ORF10
0
0.5
1
1.5
2
Alpha
Beta
Gamma
Delta
BA.1
BA.2
BA.3
BA.4
BA.5
XBB.1
XBB.2
XBB.3
XBB.4
